# LinearKnot: linear-time ensemble analysis of RNA density-2 pseudoknots under hierarchical folding

**DOI:** 10.64898/2026.11.06.764809

**Authors:** Takumi Otagaki, Kengo Sato

## Abstract

**Motivation:** A pseudoknot is an arrangement of crossing base pairs. Pseudoknots complicate structure prediction and calculations over possible structures (an ensemble). CParty sums structure weights for hierarchical pseudoknots that keep a supplied pseudoknot-free structure (scaffold). Its time is cubic in sequence length.

**Results:** LinearKnot approximates CParty with beam pruning. It combines at most two partial structures in each step and keeps the highest-ranked candidates. At fixed beam width, hairpin length limit, and scaffold search limit, partition functions, minimum-free-energy (MFE) predictions, and representative *γ*-centroid structures from base-pairing probabilities take linear time in sequence length. Loop profiles give probabilities for pairing and six unpaired loop types, including gaps between crossing stems (PseudoLoop). Window accessibility is the probability that a window is unpaired. For each RNA, we compared each base’s distributions over pairing partners (including the unpaired state) and structural contexts with exact probabilities and averaged their total variation (TV) distances over bases unpaired in the scaffold. At width 200, the 90th percentiles of these RNA averages among 300–399-base RNAs were 0.0015 for pairing and 0.0017 for contexts. At width 100, LinearKnot completed partition functions for all 107 real RNAs through 29 kilobases. Under our settings, CParty completed RNAs up to about 1 kilobase and was faster on them. For the RNAcentral long non-coding RNA URS00023A4DB9 (29,106 nt), partition-function, pairing-probability, and MFE calculations at width 100 took about 7 minutes and 3.22 GiB. We compared free energy and pairing probabilities before and after each increase in beam width and local limits. These changes generally became smaller at larger widths. At width 200, the calculations for URS00023A4DB9 took about 20 minutes and 5.68 GiB. In rat telomerase RNA, a four-base window next to a pseudoknot was mostly unpaired as part of a PseudoLoop. A pseudoknot-free calculation with the same scaffold gave lower accessibility and placed the window mainly in a hairpin when unpaired.

**Availability and implementation:** LinearKnot provides the linearknot command at https://github.com/TakumiOtagaki/LinearKnot. Experiment scripts and data are at https://github.com/TakumiOtagaki/LinearKnotExperiments.

## 1. Introduction

RNA can form different structures by changing which bases pair with each other. The structure determines which bases can interact with other molecules [Dethoff et al., 2012]. A minimum-free-energy (MFE) structure has the lowest energy among the structures considered. An ensemble describes a set of possible structures and their probabilities. Lower-energy structures have larger Boltzmann weights, and the partition function sums these weights to normalize the probabilities. A base-pairing probability is the probability that two particular bases pair. McCaskill’s algorithm computes the partition function and base-pairing probabilities for pseudoknot-free structures, which have no crossing base pairs [McCaskill, 1990]. The ViennaRNA 2.0 software package provides these calculations and other RNA structure analyses [Lorenz et al., 2011]. CapR gives loop profiles: structural-context probabilities that describe whether each base is paired or belongs to a particular loop type [Fukunaga et al., 2014]. Raccess gives window accessibility, the probability that all bases in an interval are unpaired together [Kiryu et al., 2011]. Boltzmann sampling draws complete structures according to their thermodynamic probabilities.

A pseudoknot is an arrangement of base pairs that cross when drawn as arcs above a sequence. When all pseudoknots are allowed, finding an MFE structure is NP-hard for a broad class of loop-based energy models [Lyngsø and Pedersen, 2000]. No known algorithm solves all such cases efficiently. Many methods therefore restrict the allowed pseudoknots [Akutsu, 2000]. Dirks and Pierce compute a partition function over a restricted pseudoknot class [Dirks and Pierce, 2003]. Knotty minimizes free energy over the pseudoknot class of Chen, Condon, and Jabbari, which includes pairs between two hairpin loops [Jabbari et al., 2018]. IPknot and ThreshKnot predict pseudoknotted structures from base-pairing probabilities [Sato et al., 2011, Sato and Kato, 2022, Zhang et al., 2019]. Both can use LinearPartition to handle long sequences [Zhang et al., 2020]. Hierarchical folding assumes that a pseudoknot-free structure forms first and additional pairs form between its unpaired regions. HFold predicts structures based on hierarchical folding [Jabbari et al., 2008]. It uses this first structure as a scaffold and predicts an extension of additional pairs between its unpaired bases. Iterative HFold refines an initial structure through repeated folding steps [Jabbari and Condon, 2014]. The extension is pseudoknot-free, but its pairs can form pseudoknots with scaffold pairs. HFold restricts these pseudoknots to the density-2 class described in Methods. CParty decomposes these structures into smaller regions and sums their Boltzmann weights, counting each structure once [Gray et al., 2024]. The sum is a conditional partition function because all structures retain the scaffold pairs. PRISM uses CParty’s partition-function calculation to sample structures according to their Boltzmann probabilities [Gray et al., 2026a]. From sample frequencies, PRISM estimates pairing and unpaired probabilities and predicts representative structures. PRISM also groups sampled structures that have the same connections between helices, even when their exact base-pair positions differ. CParty’s cubic running time and quadratic memory use make long-RNA analyses costly.

Beam pruning offers a way to process long sequences by limiting the partial structures retained during a left-to-right calculation. The beam width sets the number retained at each selection step. LinearFold uses this approach for MFE prediction [Huang et al., 2019]. LinearPartition approximates partition functions and base-pairing probabilities [Zhang et al., 2020]. LinearSampling draws Boltzmann samples from the approximate ensemble computed by LinearPartition [Zhang et al., 2023], and LinearCapR computes structural-context probabilities [Otagaki et al., 2026]. These methods analyze pseudoknot-free structures. CapR and Raccess also exclude pseudoknots, so these loop-profile and accessibility methods omit unpaired regions formed by crossing pairs. For hierarchical folding with pseudoknots, Spark omits computations that cannot improve HFold’s MFE optimum (sparsification) [Gray et al., 2026b]. This saves time and memory but does not give a linear-time bound.

We present LinearKnot for structure prediction and ensemble analysis with pseudoknots under a supplied scaffold. We reorganize CParty’s decomposition for beam pruning, giving linear time and space in sequence length at fixed beam width and local limits. LinearKnot combines partial structures two at a time and prunes candidates at each right endpoint. The calculation sums Boltzmann weights for the partition function or minimizes energy for MFE prediction. LinearKnot computes pairing and unpaired probabilities analytically in the ensemble kept by beam pruning, whereas PRISM estimates them from sample frequencies. From the partition function, LinearKnot computes base-pairing probabilities, loop profiles, window accessibility by loop type, Boltzmann samples, and representative *γ*-centroid structures chosen from base-pairing probabilities. The loop profiles include PseudoLoop, the unpaired gaps between the crossing stems of a pseudoknot. We validate the calculations, measure approximation errors and resource use, and illustrate the ensemble analyses with telomerase RNA.

## 2. Methods

### 2.1. Input, conditional ensemble, and energy

Let *S* be an RNA sequence with *n* nucleotides (nt), numbered 1 to *n*. A structure is a set of base pairs (*i, j*) with *i < j*; each base participates in at most one pair. Pairs (*i, j*) and (*k, l*) cross when *i < k < j < l* or *k < i < l < j*. The user supplies a pseudoknot-free secondary structure, the scaffold *G*. Section 2.5 describes how we construct scaffolds for the experiments. We retain every pair in *G* and add a pseudoknot-free extension *G*′ between bases unpaired in *G*. We call these bases free bases. The completed structure is *R* = *G* ∪ *G*′ (Figure 1A). Pairs in the extension *G*′ can be AU, UA, CG, GC, GU, or UG. A hairpin is an unpaired segment closed by a base pair and must contain at least three unpaired bases.

**Fig. 1:**
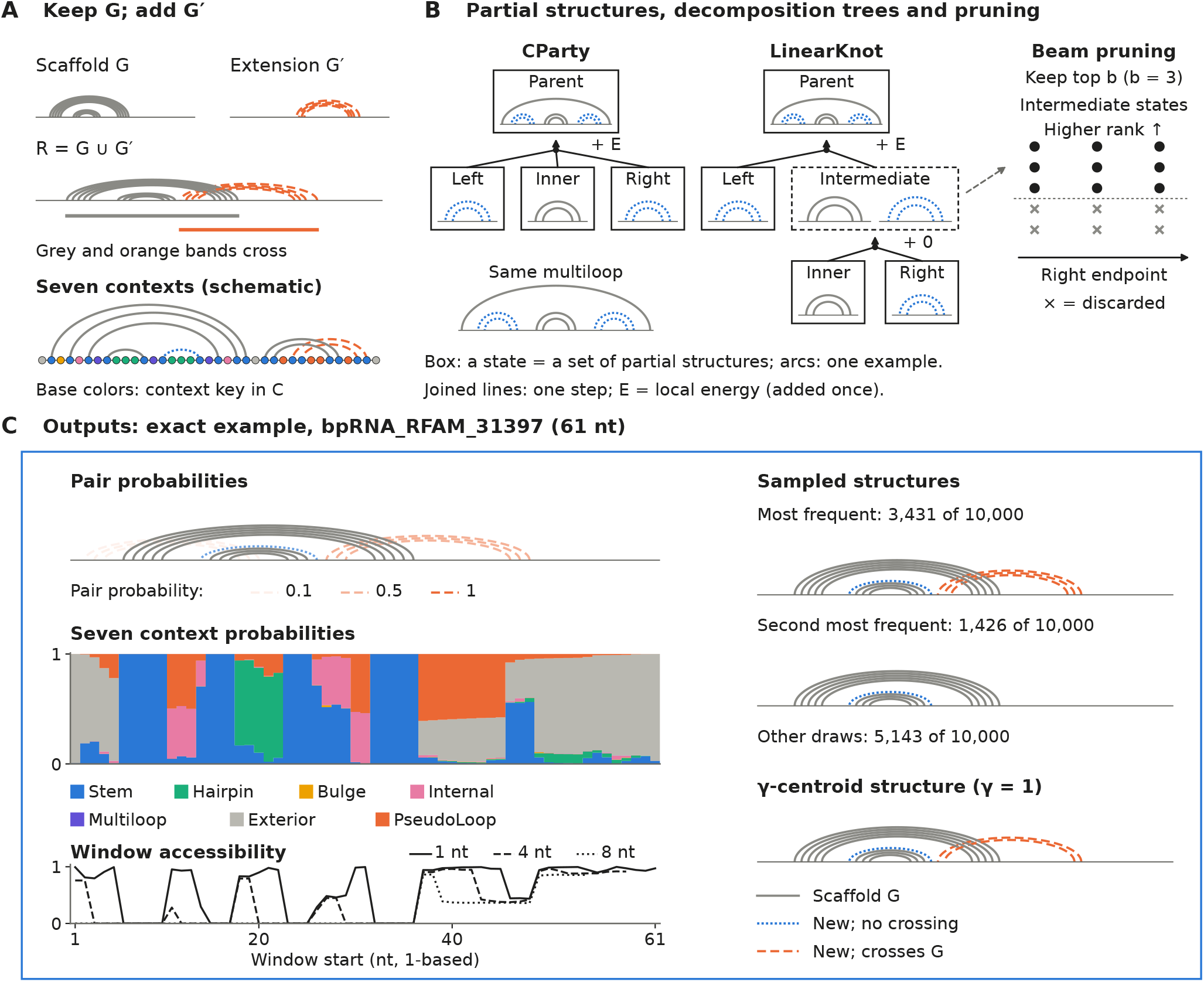
Structure prediction and conditional ensemble analysis with LinearKnot. **(A)** Scaffold *G* and its extension each have no crossing pairs and share no paired bases. Their union forms *R*. The grey and orange underlines mark the spans of two bands whose pairs cross each other. Density-2 allows at most two bands in a connected crossing group to span one position. The lower sketch in A shows a schematic structure with each base colored by its structural context (key in C). Orange unpaired bases lie in gaps between crossing bands (PseudoLoop). **(B)** A multiloop connects closed substructures called branches. Between two pairs of one scaffold band, CParty combines a multiloop’s inner part with both new side branches in one step. CParty already combines parts of ordinary multiloops two at a time. Boxes represent states (sets of partial structures); arcs show examples. LinearKnot first joins the inner part and right branch (dashed box), then adds the left branch. Only the second step adds local energy *E*. Pruning includes intermediate states. The right endpoint is a partial structure’s last sequence position. The main beam keeps the top *b* states per type and endpoint (*b* = 3). Circles mark retained candidates; crosses mark discarded candidates. The same decomposition gives partition functions by summing weights and MFE structures by minimizing energy. **(C)** An exact calculation for bpRNA RFAM 31397 (61 nt) gives outputs from one conditional ensemble. Its reference contains a pseudoknot; *G* retains only the eight AU, GC, or GU pairs in its nested part. Solid grey arcs show *G*; dotted blue arcs show new non-crossing pairs; dashed orange arcs show new pairs crossing *G*. In the upper-left plot, arc opacity shows pair probability (at least 0.05). Each other arc diagram shows one sampled or *γ*-centroid structure. The profile shows seven context probabilities (Methods). Accessibility is the probability that all window bases are unpaired. Accessibility curves use solid lines for 1-nt windows, dashed lines for 4-nt windows, and dotted lines for 8-nt windows. Window starts are 1-based; position 20 on the 8-nt curve represents bases 20–27. The two sample diagrams show the most frequent structures (3,431 and 1,426 of 10,000 draws). The representative structure uses *γ* = 1.

Following HFold and CParty, we restrict completed structures to the density-2 class [Jabbari et al., 2008, Gray et al., 2024]. Nested pairs have both endpoints of one pair between those of the other. A band is the set of all nested pairs that cross exactly the same nonempty set of other pairs. A band’s span runs between its outermost pair’s endpoints. The density-2 condition requires that, in each group of bands connected directly or indirectly by crossings, at most two bands span any position. For sequence *S* and scaffold *G*, let Ω(*S, G*) be the set of completed structures allowed by these conditions and the decomposition rules in Supplementary Sections S1.1 and S1.3. An internal loop is an unpaired region between two nested base pairs. We allow at most 30 unpaired bases in an internal loop closed by a new non-crossing pair and fewer than 30 for a new crossing pair (Supplementary Section S1.4). Exact calculations sum over Ω(*S, G*) without beam pruning and without the hairpin-length and scaffold-depth limits defined below.

Ordinary loops use Turner 2004 parameters; pseudoknots use CParty’s constants from the re-estimated Dirks–Pierce set (DP09) [Mathews et al., 2004, Andronescu et al., 2010, Gray et al., 2024]. LinearKnot can also use DP09 parameters for ordinary loops. The energy *E*_*G*_(*R*) is the sum of local loop terms assigned by the decomposition. Let *T* be absolute temperature, *R*_gas_ the gas constant, and *κ* = *R*_gas_*T* . The Boltzmann weight *w*_*G*_(*R*), conditional partition function *Z*, and structure probability are

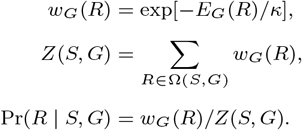

All experiments use 37 °C. Supplementary Section S1.2 specifies the local energy settings.

### 2.2. Reorganizing CParty’s decomposition for beam pruning

CParty decomposes each allowed structure into smaller parts in just one way [Gray et al., 2024]. This decomposition forms a tree: each step joins smaller parts and adds a local energy term. The trees share parts, forming a directed hypergraph whose edges connect child parts to their parent. Starting with the smallest parts, the inside calculation sums Boltzmann weights over these trees. Each step sums products of its local Boltzmann factor and child inside weights. The sum for the full sequence is *Z*. Weighted decompositions also give base-pairing and structural-context probabilities, window accessibility, Boltzmann samples, and MFE and *γ*-centroid structures. Computing these quantities from the decomposition follows Ponty and Saule’s general approach [Ponty and Saule, 2011].

CParty already combines parts of ordinary multiloops two at a time. For a multiloop between two pairs of one scaffold band, CParty joins the inner part and both side branches in one step (Figure 1B). LinearKnot combines partial structures two at a time, so it can prune before making all combinations. With beam width *b*, the main selection keeps the top *b* comparable candidates per group at each right endpoint. Supplementary Sections S1.3 and S1.4 specify the combinations and candidate groups.

LinearPartition ranks each interval by its inside weight times the exterior-loop weight for the sequence to its left [Zhang et al., 2020]. There, each part covers one interval, and child intervals join end to end. Here, some parts cover two separated intervals, such as a band’s paired segments, so the left sequence alone does not estimate their surroundings. We therefore run a small-beam preliminary calculation and use it to estimate their surrounding weights for ranking (Supplementary Section S1.4).

Candidate generation also uses two local limits. The first, *h*, bounds the unpaired length of a new hairpin. The second, *d*, limits how far we search through scaffold loops for a new pair that crosses *G*. Starting at the innermost scaffold pair enclosing a free base, we follow the enclosing loops outward. We count only loops with at least two scaffold branches, up to *d* such levels. The starting pair’s loop is not counted; the exterior loop counts if it has at least two branches. Stacked pairs and internal loops do not count toward this depth. We use *h* = *d* = max(*b*, 30). We counted these levels in the predicted scaffolds used for the bpRNA and RNAcentral experiments, rather than in reference structures. These scaffolds were built from LinearPartition pairing probabilities at beam width 200, retaining pairs with probability at least 0.7 in helices of at least two stacked pairs (Section 2.5). The counts were at most 5 in bpRNA below 400 nt, 16 in longer bpRNA, and 28 in the seven RNAcentral long noncoding RNAs. With these predicted scaffolds, no search reached *d* = 30 (Supplementary Table S4). This is an observation for these scaffolds, not a general bound; scaffolds with deeper branching, such as those from other prediction rules or reference structures, may need a larger *d*, which users can set. Exact calculations use neither limit, but both exact and finite-beam modes retain the internal-loop restrictions in Supplementary Section S1.4. Let 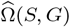 be the set of completed structures that remain after pruning and the local limits. Their total weight is 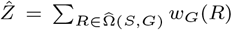, and their probabilities are 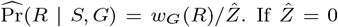, no completed structure remains and we leave probabilities undefined.

### 2.3 Probabilities, sampling, and representative structures

After pruning, we keep selected parts and their decomposition steps. Outside traverses these steps in reverse and sums the weights of surrounding structures. The inside weight times the outside weight, divided by 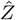, gives the probability that a structure contains that part. Supplementary Section S1.5.1 gives the equations. Pairing, context, accessibility, sampling, and *γ*-centroid calculations reuse the partition-function calculation without further pruning. Let *p*_*ij*_ be the base-pairing probability (BPP) of positions *i* and *j*, with *p*_*ji*_ = *p*_*ij*_ . The unpaired probability is *q*_*i*_ = 1 − ∑_*j*≠ *i*_ *p*_*ij*_, where the sum covers all other sequence positions. Every scaffold pair (*i, j*) ∈ *G* has base-pairing probability *p*_*ij*_ = 1.

We assign each base to one of seven structural contexts (Figure 1A, lower sketch). Stem denotes paired bases. Hairpin denotes unpaired bases at the end of a stem, a run of consecutive nested pairs. Bulge denotes unpaired bases on one side between nested pairs; Internal denotes unpaired bases on both sides.

Multiloop denotes unpaired bases in a loop with at least two child branches. Exterior denotes unpaired bases outside closed regions. PseudoLoop denotes unpaired bases in the gaps between crossing bands (Figure 1C). For context label *c*, let *P*_*i*_(*c*) be the probability that base *i* has that context. These seven events are mutually exclusive. Thus, *P*_*i*_(Stem) = 1 − *q*_*i*_, and the six unpaired-context probabilities sum to *q*_*i*_. Supplementary Section S1.5.2 gives the classification boundaries and aggregation checks.

For a window of length *w* starting at position *s*, window accessibility *A*(*s, w*) is the probability that all its bases are unpaired together. Here, 1 ≤ *w* ≤ *n* and 1 ≤ *s* ≤ *n* − *w* + 1. Let end(*R*) be the positions paired in structure *R*. Then

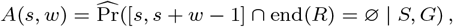

where ∅ is the empty set. The interval [*s, s* + *w* − 1] lists the window positions. Accessibility is a joint probability, rather than a product of individual *q*_*i*_ values. We separate accessibility into six contributions, one for each unpaired loop type. Each component is the joint probability that all window bases are unpaired and share that type. The six components sum to *A*(*s, w*). Supplementary Section S1.5.3 gives the calculation. A window containing a scaffold-paired base has accessibility 0, and *A*(*s*, 1) = *q*_*s*_. The window’s opening free energy is −*κ* log *A*(*s, w*), or +∞ when accessibility is zero.

Boltzmann sampling starts from the full sequence and chooses each decomposition step in proportion to its local weight times the inside weights of the smaller parts joined in that step. Repeating this choice through the retained graph draws structures with probability 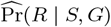. For a representative structure, we use the *γ*-centroid estimator with BPPs from the same ensemble [Hamada et al., 2009]. For a candidate structure 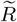 and a nonnegative pair weight *γ*, it maximizes

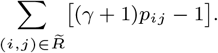

Larger *γ* gives more weight to correctly predicted pairs. We select 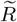 from the retained ensemble without further pruning. We use *γ* = 1 in the examples.

Minimum-free-energy prediction uses the same decomposition but ranks candidates by best-energy scores and prunes separately. At finite beam width, it can retain different candidates from the partition-function calculation and minimizes energy over its own retained structures.

### 2.4. Computational complexity

Assume a supplied scaffold and 2 ≤ b ≤ n. Fix the local limits h and d and the number of window lengths. The partition function, BPPs, loop profiles, window accessibility, and MFE and γ-centroid predictions use expected time T_calc_ and space M :

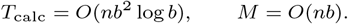

At fixed beam width these costs are linear in sequence length. The *O*(*nb*) space bound excludes sampling. Supplementary Section S2 gives the assumptions, sampling costs, and dependence on the hairpin-length and scaffold-search limits.

### 2.5. Experimental settings

Unless stated otherwise, we construct experimental scaffolds from LinearPartition pairing probabilities at beam width 200. We select all pairs with probability at least 0.7, then retain only helices of at least two stacked pairs. Numerical validation compares compatible partition functions with CParty and checks our ensemble by complete enumeration on small inputs. Accuracy evaluation uses 26,063 bpRNA-1m(90) sequences of at least 50 nt with only A, C, G, and U [Danaee et al., 2018]. We compare *b* = 20, 50, 100, 200 with exact calculations below 400 nt. For longer sequences, we compare these widths and *b* = 300 with a finite reference at *b* = 400.

Total variation (TV) distance measures finite-beam probability error. For distributions *P, Q* on a set *X*, 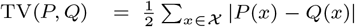. For 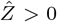,let 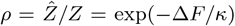 be the fraction of the exact weight that remains, where 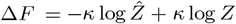. Suppose 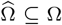 with unchanged weights, and write Pr(*R*) for Pr(*R* | *S, G*). Then remaining structures have probability Pr(*R*)*/ρ* after pruning, removed ones have probability 0, and

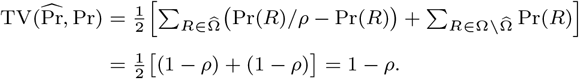

The two sums are the probability gained by retained structures and the probability lost with removed structures. The pairing or context distribution at a base sums structure probabilities in groups. Its TV and each window’s absolute accessibility error are at most 1 − *ρ* (Supplementary Section S1.5.4).

At each free base, we compute TV over pairing partners and the unpaired state, or over the seven contexts. We average over free bases within each RNA, then report percentiles across RNAs. For accessibility, we average absolute probability differences over windows containing only free bases within each RNA, then summarize across RNAs. RNAs without eligible windows are omitted from that metric.

Resource measurements use seven RNAcentral sequences and synthetic sequence–scaffold controls [RNAcentral Consortium, 2021]. We use beam width *b* = 100, limit new hairpins to 100 unpaired bases, and search at most 100 enclosing scaffold multiloops. We measured each complete command’s wall-clock (elapsed) time and peak memory (maximum resident set size). Values are medians of three runs after one warm-up, excluding scaffold construction. TSUBAME4.0 at Institute of Science Tokyo has two 96-core AMD EPYC 9654 processors (2.4 GHz) per node. Each 40-core allocation ran up to ten single-threaded processes (Supplementary Section S3.3). CParty and Spark comparisons used 107 real RNAs on 16-core allocations of TSUBAME4.0. Partition-function settings matched between CParty and LinearKnot (Supplementary Section S3.6).

We use rat telomerase RNA (Terc, 388 nt; NR 001567.1) to illustrate the ensemble analyses. Its scaffold contains the 61 AU, GC, or GU pairs in the nested part of the bpRNA RFAM 4317 reference, with *j* − *i >* 3. We compare *b* = 200 with exact calculations. We show four-base accessibility by loop type, draw 1,000 samples, and predict the *γ*-centroid structure at *γ* = 1.

## 3. Results

### 3.1. Validation and approximation accuracy

We compared partition functions with a corrected version of CParty. The maximum absolute log *Z* difference was 1.9 × 10^−12^ on 305 inputs (Supplementary Section S1.7). Enumeration of small structures checked the structure weights, pairing and context probabilities, accessibility, MFE energies, and sampling frequencies (Supplementary Section S3.1). With an empty scaffold, log *Z* and BPPs agreed with ViennaRNA within 1.5 × 10^−12^. For small inputs, *γ*-centroid scores agreed with enumerated maxima for each retained ensemble within 2 × 10^−13^ (Supplementary Section S3.1).

We measured beam-width effects on ensemble weights, pairing probabilities, loop profiles, and total window accessibility (Figure 2). For 24,625 RNAs below 400 nt, we used exact references. Larger beam widths usually reduced the differences. At *b* = 200, absolute free-energy differences were below 7 × 10^−13^ kcal/mol for tested RNAs below 200 nt and below 0.012 for at least 90% of 200–399-nt RNAs.

**Fig. 2:**
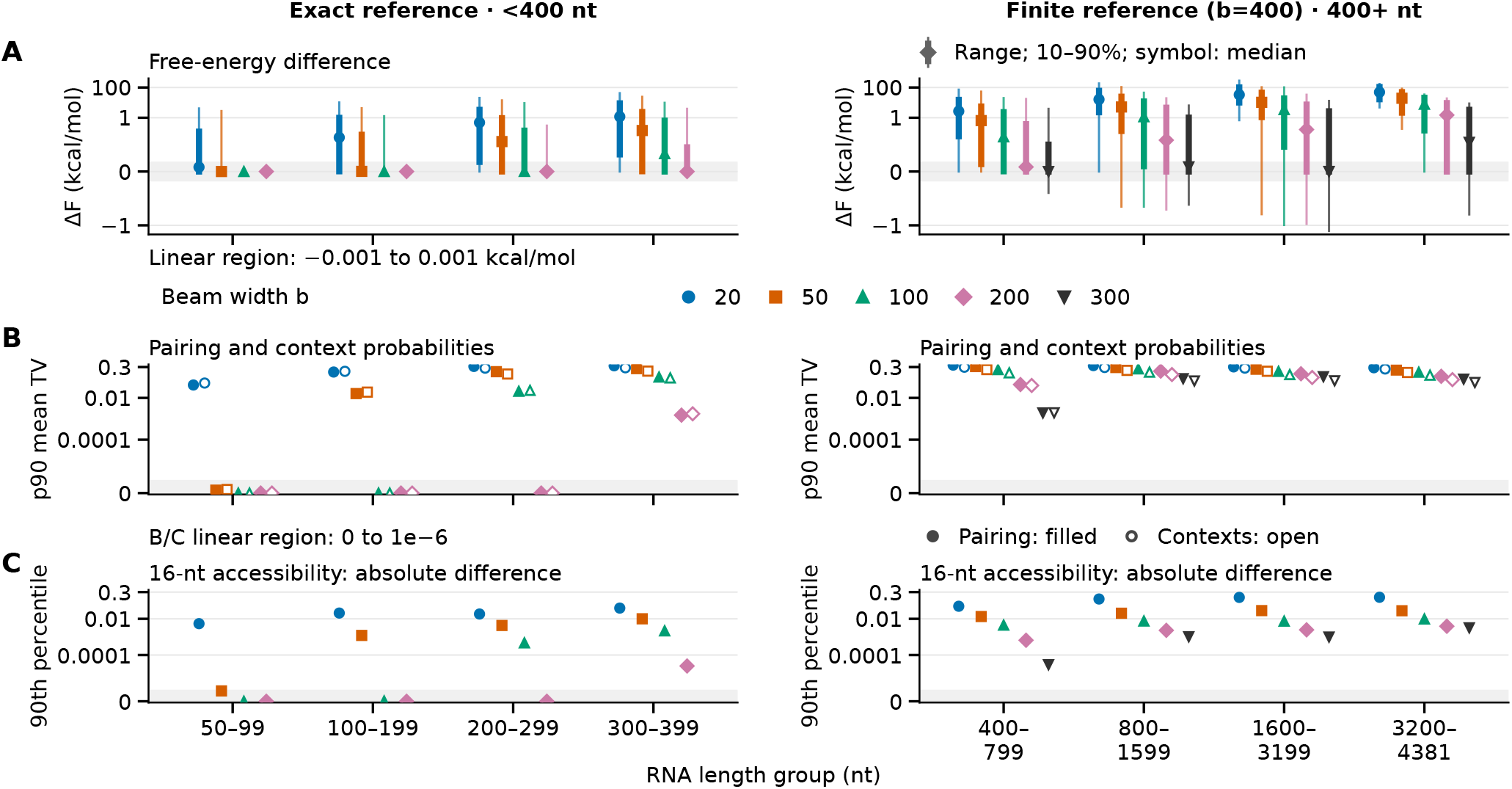
Sensitivity of conditional-ensemble outputs to beam width and local limits. Left panels use exact references for 24,625 RNAs below 400 nt; right panels use *b* = 400 for 1,438 RNAs of 400–4,381 nt. Colors and shapes mark beam width *b* throughout, with beam widths equally spaced within each length group. Local limits are *h* = *d* = max(*b*, 30). The *b* = 400 self-comparison is omitted. **(A)** The signed difference ∆*F* is the finite-beam ensemble free energy minus the reference free energy. Thin lines show full ranges; thick lines show the 10th–90th percentiles; symbols show medians across RNAs. The shaded band is linear between −0.001 and 0.001 kcal/mol; the scale is logarithmic beyond it. **(B)** Filled symbols compare distributions over pairing partners and the unpaired state; open symbols compare seven context probabilities. Total variation (TV) distance is half the sum of absolute probability differences. We average TV over bases unpaired in *G* within each RNA, then take the 90th percentile across RNAs in each length group. **(C)** For each RNA, we average the absolute accessibility differences over 16-nt windows containing only bases unpaired in *G*, including windows with zero accessibility. Points show the 90th percentile of these averages across RNAs. B and C are linear from 0 to 0.000001 (shaded) and logarithmic above it; zeros remain zero. Supplementary Figure S3 shows 1-, 4-, 8-, and 16-nt windows.

Among the 1,095 RNAs of 300–399 nt, the 90th percentile, or p90, of ∆*F* fell from 8.94 kcal/mol at *b* = 20 to 0.0111 at *b* = 200. At *b* = 200, p90 mean TV was 0.0015 for BPPs and 0.0017 for contexts. Of 1,048 RNAs with eligible 16-nt windows, at least 90% had mean absolute accessibility error at most 2.5 × 10^−5^. Maximum mean TV was 0.37 for BPPs and 0.21 for contexts.

For 1,438 RNAs of 400–4,381 nt without exact references, Figure 2B and Supplementary Table S5 show sensitivity to beam width and local limits.

On seven RNAcentral inputs, changes between successive widths generally became smaller as beam width and local limits increased (Supplementary Figure S5). For *b* = 100 → 200, 200 → 300, and 300 → 400, maximum absolute free-energy changes were 0.00463, 0.00126, and 0.000507 kcal/mol per base. We computed TV between the distributions over pairing partners and the unpaired state, then averaged over free bases within each RNA. The largest RNA averages were 0.1518, 0.0581, and 0.0468 for these three increases, respectively. URS000238C72C (22,543 nt) still had maximum per-base TV 0.9991 between *b* = 300 and 400. The 29,106-nt RNA took about 20 min and 5.68 GiB at *b* = 200 in one measurement.

### 3.2 Loop profiles and accessibility by loop type around a pseudoknot

We examined a predicted accessible pseudoknot-loop window in Terc (Figure 3). Comparative studies provide a vertebrate telomerase RNA secondary-structure model [Chen et al., 2000]. The scaffold retains 61 canonical nested pairs from the bpRNA RFAM 4317 reference [Danaee et al., 2018]. The six outer reference pairs (O) crossing *G* are (63, 139), (64, 138), …, (68, 134).

**Fig. 3:**
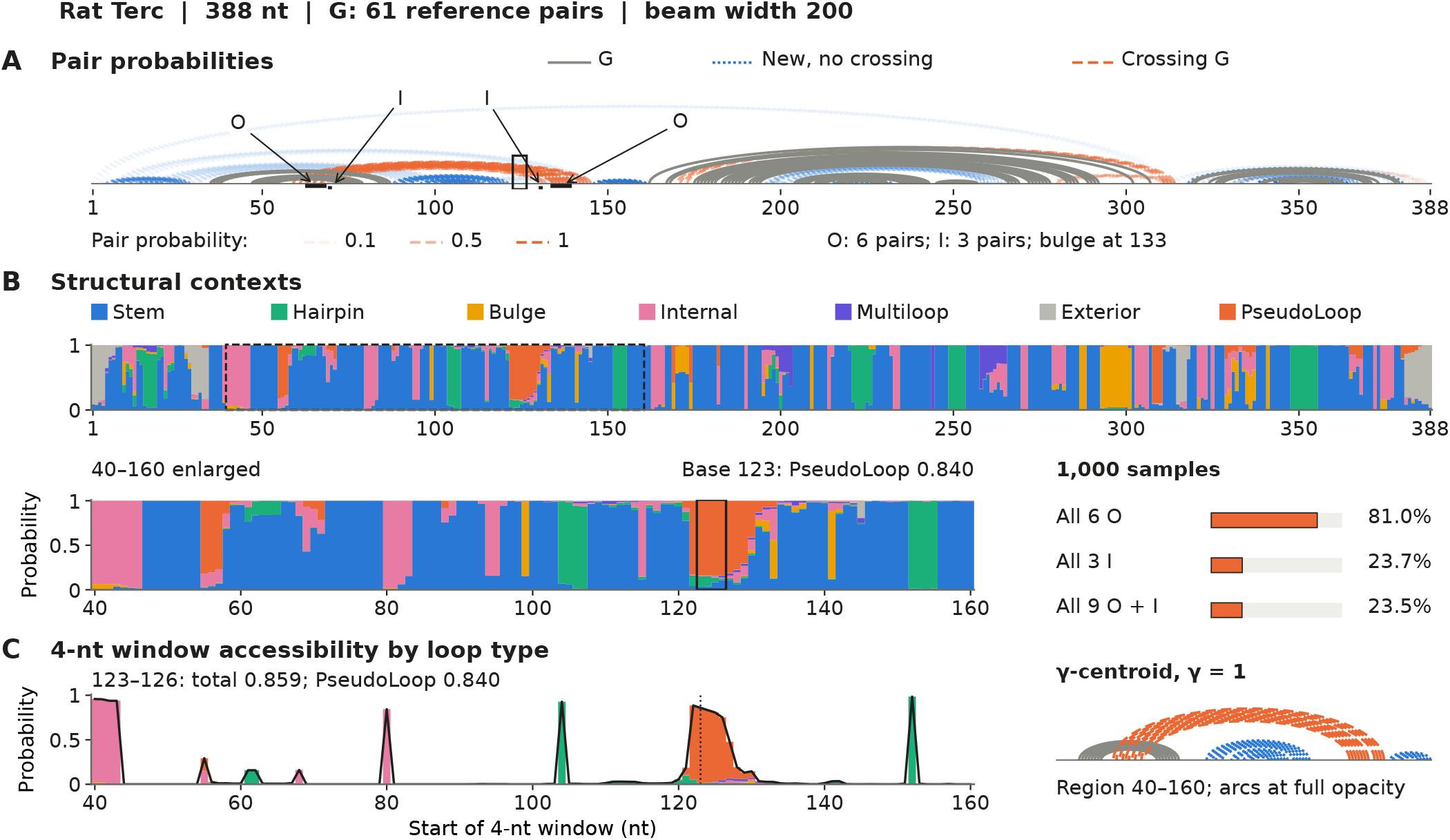
Loop profiles and accessibility by loop type in rat Terc (NR 001567.1; 388 nt) at beam width 200. The scaffold *G* contains 61 canonical AU, GC, or GU pairs from the nested part of the bpRNA reference. Within coordinates 40–160, maximum absolute differences from exact calculations are below 0.00284 for contexts and 0.00192 for four-base total accessibility. **(A)** Arcs show pairs with probability at least 0.05. Grey solid arcs show *G*; blue dotted arcs show new non-crossing pairs; orange dashed arcs show new pairs crossing *G*. Arc opacity shows pair probability, as illustrated by the numeric key. O marks six outer reference pairs, (63, 139), (64, 138), …, (68, 134). I marks three inner reference pairs, (69, 132), (70, 131), (71, 130). Bulged base 133 separates the two groups. Together they form the nine reference pairs that cross *G* in this region. The outlined region marks bases 123–126. **(B)** Colored bands show seven single-base context probabilities. PseudoLoop is the context of unpaired gaps between crossing bands, defined in Methods. The enlarged track and C share coordinates 40–160. The outlined region spans bases 123–126; the guide in C marks window start 123. **(C)** Each colored component is the joint probability that all four window bases are unpaired and assigned to that loop type. The black line is their sum. Window 123–126 has accessibility 0.859, including a PseudoLoop contribution of 0.840. The small bars at right show fractions of 1,000 samples: all six O pairs in 810 draws, all three I pairs in 237, and all nine pairs in 235. Other pairs are allowed, and these frequencies have sampling uncertainty. Below the bars, the *γ*-centroid structure at *γ* = 1 shows region 40–160 with fully opaque arcs. Accessibility coordinates denote window starts.

The three inner reference pairs (I) are (69, 132), (70, 131), (71, 130). Bulged base 133 separates these two groups (Figure 3A).

At *b* = 200, probabilities were 0.83–0.84 for the six O pairs and 0.28–0.34 for the three I pairs. About 98% of the unpaired probability mass of window 123–126 belongs to PseudoLoop. Under this scaffold and energy model, this window is therefore likely unpaired, and its unpaired states fall almost entirely in a pseudoknot loop. Experiments must test whether this window is physically exposed or binds other molecules.

At *b* = 200 and in exact mode, *γ*-centroid structures with *γ* = 1 contained all six O pairs, (63, 139), …, (68, 134), and none of the I pairs, (69, 132), (70, 131), (71, 130). All nine reference pairs occurred together in 235 of 1,000 Boltzmann samples (23.5%). MFE structures at *b* = 200 and in exact mode contained all nine reference pairs. New pairs crossing *G* were predicted near the 3′ end but absent from the reference. Pair probabilities describe one pair at a time. The 23.5% sample fraction estimates the probability that all nine reference pairs occur together under this scaffold and energy model.

We compared the exact LinearKnot ensemble, where new pairs may cross *G*, with pseudoknot-free structures, where new pairs cannot cross *G*. For the pseudoknot-free comparison, ViennaRNA 2.7.2 enforced the same 61 scaffold pairs as hard constraints and computed window accessibility as the partition function with bases 123–126 additionally forced unpaired divided by the partition function with only the scaffold constraints. Exact accessibility of window 123–126 was 0.861 with LinearKnot and 0.123 in the pseudoknot-free comparison. The PseudoLoop contribution was 0.842 with LinearKnot; the Hairpin contribution was 0.117 in the pseudoknot-free comparison. In the pseudoknot-free comparison, the window has lower accessibility and is mainly in a hairpin when unpaired.

### 3.3 Time, memory, and analyses of long RNAs

For URS00023A4DB9 (29,106 nt), partition function, BPP, and MFE calculations took about 7 min and 3.22 GiB (Figure 4A and B). Loop profiles took about 5 min and 3.0 GiB with or without total accessibility (Supplementary Section S3.3).

**Fig. 4:**
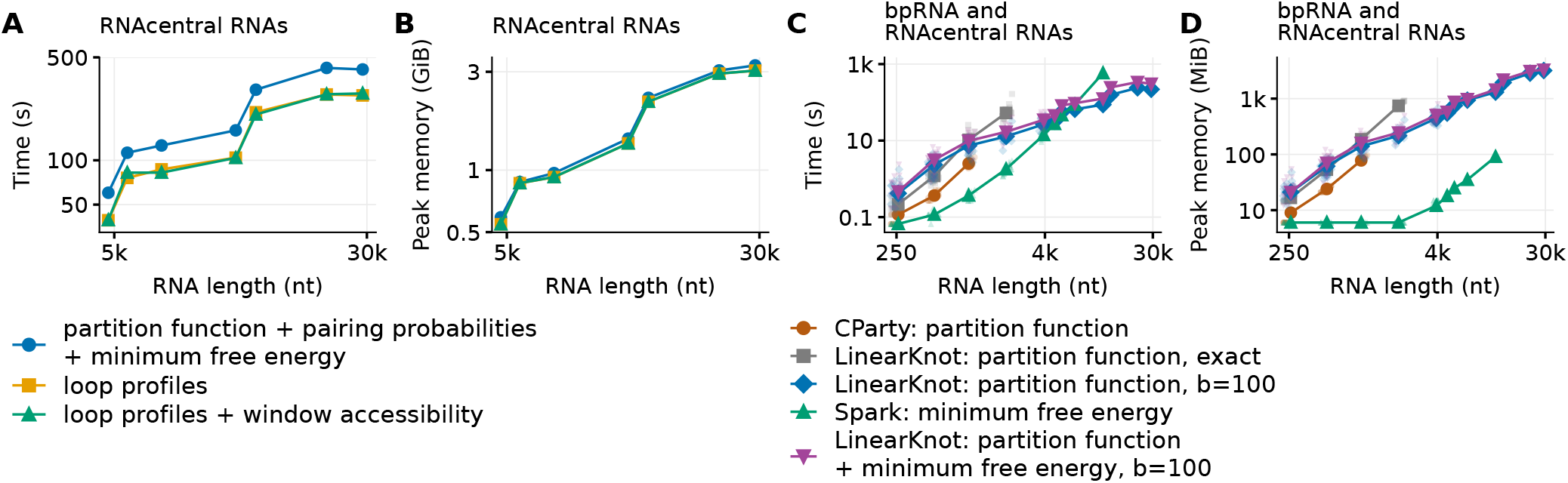
Wall-clock time and peak memory; logarithmic axes. **(A, B)** Seven RNAcentral inputs, *b* = *h* = *d* = 100. Calculations: partition function, BPPs, and MFE; loop profiles; loop profiles with total window accessibility (1-, 4-, 8-, 16-nt windows). Points show medians of three measurements after one warm-up. **(C, D)** 100 bpRNA inputs (20 per group) and seven RNAcentral inputs. Partition function comparisons use the same energy and boundary settings. Spark: MFE; purple LinearKnot: partition function and MFE, no BPP (*b* = *h* = *d* = 100). Faint points: input medians; lines: group medians or individual RNAcentral inputs. CParty completed no input in the groups of 2,000 nt or longer and failed on one 994-nt input in the 1,000-nt group. Most failures were numerical overflows when run without scaling because of CParty’s scaling behavior; some inputs exceeded the time limit.

Under matched partition-function settings, CParty was faster on all 59 RNAs completed by both methods (225–1,077 nt; Figure 4C and D). Its median peak memory was 9, 24, and 78 MiB in the 250-, 500-, and 1,000-nt groups. CParty completed 19 of 20 inputs in the 1,000-nt group, with numerical overflow on the 994-nt input, and none in the groups of 2,000 nt or longer. Of its 48 failed inputs, 43 overflowed when run without scaling because of CParty’s scaling behavior. Some longer inputs exceeded the time limit; others crashed or reached the memory limit (Supplementary Section S3.6). These failures prevented a speed comparison on longer inputs. At 11,832 nt, LinearKnot took 126 s and 1,450 MiB versus Spark’s 581 s and 91 MiB. The comparison measures resources: Spark computes MFE; LinearKnot computes partition function and MFE (Supplementary Section S3.6).

Supplementary Figure S4 shows the synthetic costs.

At *b* = 100, the six analyses listed in Supplementary Table S8 also completed for 22,743-nt NEAT1 2.

## 4. Discussion

LinearKnot computes probabilities over the ensemble of structures that keep a supplied scaffold and may add pairs crossing it.

Crossing pairs therefore affect every output, not only their own pairing probabilities. In Terc, for example, window 123–126 was unpaired more often, and mainly inside a PseudoLoop rather than a hairpin, once crossing pairs were allowed. Inside and outside calculations give these probabilities without sampling error. Boltzmann samples additionally show which pairs occur together. Finite-beam MFE prediction prunes separately and may keep different structures from the probability calculations.

Adjustable beam width lets users balance time and memory costs against changes in their estimates, an advantage over exact CParty. Larger beams usually improved agreement with exact values in short RNAs. For long RNAs, exact references were unavailable, so the comparisons between beam widths measured sensitivity to the approximation settings. Free energy per base changed little above *b* = 100, while pairing distributions continued to change as beam width increased (Supplementary Figure S5).

Thus, the effect of beam width depends on which quantity is being examined.

Changing the scaffold structure changes the allowed completed structures and the assignment of their energy terms. The reported probabilities therefore depend on the supplied scaffold as well as the RNA sequence. Our experimental scaffolds use a minimum LinearPartition pair probability of 0.7 and helices of at least two stacked pairs. Users can change this probability threshold or supply a scaffold based on other structural information.

Agreement with reference structures also depends on the energies assigned to the allowed structures. We examined this dependence by predicting structures with scaffolds taken from the nested parts of reference structures (Supplementary Section S3.5). Even exact predictions included few reference pseudoknot pairs. Because these calculations used no beam pruning, pruning alone cannot explain the low agreement. Using the DP09 parameter set instead of Turner 2004 for ordinary loops also changed which reference pairs were predicted. These results show that evaluating pseudoknot predictions requires attention to the energy model as well as the scaffold and beam width.

## 5. Conclusion

LinearKnot analyzes density-2 pseudoknot ensembles under hierarchical folding for RNAs tens of thousands of bases long. For a supplied scaffold, its probabilities, samples, and *γ*-centroid structures from one retained ensemble show where crossing pairs and open windows are likely.

## 6. Data availability

LinearKnot (version 0.2.2) is available at https://github.com/TakumiOtagaki/LinearKnot. This version is archived at Zenodo, https://doi.org/10.5281/zenodo.24006507. Experiment scripts and data are at https://github.com/TakumiOtagaki/LinearKnotExperiments. The results in this paper were computed with version 0.2.1. Version 0.2.2 fixes only the probability of a user-specified complete structure, which this paper does not use.

## 7. Acknowledgements

We used the TSUBAME4.0 supercomputer at Institute of Science Tokyo for the computations in this study.

