## Supplementary Information for "LinearKnot: linear-time ensemble analysis of RNA density-2 pseudoknots under hierarchical folding"

### S1 Supplementary methods

Notation and abbreviations follow Methods, including $Z$ (exact partition function) and $\hat{Z}$ (after beam pruning).

#### S1.1 States and structural regions

A state groups partial structures by type and endpoint coordinates (Table S1). An interval is weakly closed in $G$ if no scaffold pair connects a base inside it to a base outside [4]. This condition concerns pairs in $G$ only, not pairs in $G'$. A completed pseudoknot region is the interval from the first to the last base of bands connected by crossings, together with the structures in their gaps. This interval must be weakly closed in $G$.

A branch is a closed substructure attached to a loop. An ordinary branch is closed by a nested pair. A pseudoknot branch is the whole pseudoknot region, including its crossing bands and gap structures. A pseudoknot branch can attach to an exterior loop (PS), an ordinary or band-spanning multiloop (PSM), or a gap in another pseudoknot (PSP; Table S2). V and WMB represent ordinary and pseudoknot branches, respectively.

A band-spanning multiloop is the loop bounded by outer and continuing inner pairs of one band and at least one side branch (Figure S1). The closing pairs may belong to $G$ or $G'$; pairs in the other layer cross both. Each side branch may be ordinary or a pseudoknot branch. Without side branches, the region between the two pairs forms a stack, bulge, or internal loop.

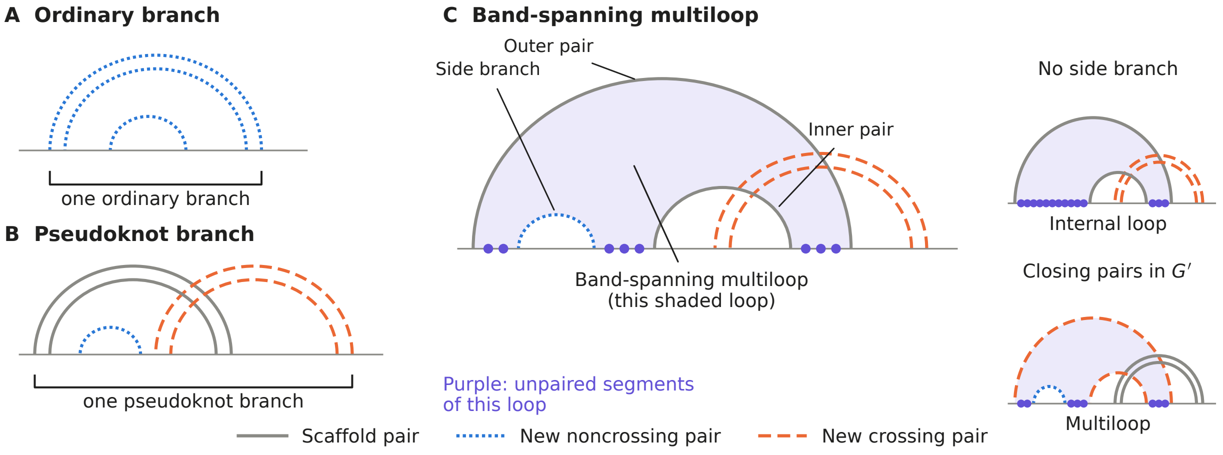


Figure S1: Branches and loops in schematic arc diagrams. A: an ordinary branch; B: a whole pseudoknot branch, including its gaps. Underlines mark one branch. C: the shaded band-spanning multiloop has a side branch and closing pairs in $G$ (main example) or $G'$ (lower inset). The upper inset shows an internal loop without a side branch. Purple dots mark unpaired segments of each shaded loop. As in Figure 1, grey solid arcs show $G$, blue dotted arcs new noncrossing pairs, and orange dashed arcs new pairs crossing $G$.

Table S1: States and structural regions. New pairs join bases unpaired in $G$, using the pair types defined in Methods.

| State | Structural region |
|---|---|
| W | Exterior region; the root is $\mathrm{W}(1, n)$. |
| V | Weakly closed region with a scaffold or allowed new closing pair. |
| VM | Ordinary-multiloop interior, including its closing energy. |
| WMv, WMp | WMv: last ordinary branch; WMp: last pseudoknot branch. Both include unpaired bases after that branch. |
| WM | Ordinary-multiloop branch sequence, with at least one branch. |
| WI, WIP | WI: weakly closed gap in a pseudoloop, with or without branches. WIP: weakly closed gap in a band-spanning multiloop, requiring a branch. |
| VP | New crossing pair and its interior, excluding the crossed scaffold-band segments. |
| VPL, VPR | VPL: crossing-pair region with a left unpaired prefix; VPR: crossing-pair region with a right extension. |
| WMBP, WMBW | Band chain not yet closed as a completed pseudoknot region; WMBW ends with a nonempty WI. |
| WMB | Completed pseudoknot region, weakly closed in $G$. |
| BE | Band segments between outer and inner scaffold pairs; excludes the inner pair's interior. |

#### S1.2 Local energy model

We use the main-text energy model: Turner 2004 or full DP09 for ordinary loops, and CParty's DP09 pseudoknot constants (Table S2) [1, 4]. For an energy contribution $x$ in kcal/mol, define $B(x) := \exp(-x/\kappa)$, using $\kappa = R_{\text{gas}}T$ from Methods. Temperature is $T = 310.15$ K. The setting for energies of unpaired bases next to helices (`dangles`) is 2.

Table S2: CParty pseudoknot constants from DP09 at 37 °C (kcal/mol).

| Constant | Value | Assigned contribution |
|---|---|---|
| PS | −1.38 | Introduce a pseudoknot region into an exterior loop. |
| PSM | 10.07 | Introduce a pseudoknot branch into an ordinary or band-spanning multiloop. |
| PSP | 15.00 | Introduce a pseudoknot branch into a pseudoloop. |
| PB | 2.46 | Add one band. |
| PUP | 0.06 | One unpaired base in a pseudoloop gap. |
| PPS | 0.96 | One closed branch in a pseudoloop. |
| $a_p, b_p, c_p$ | 3.41, 0.56, 0.12 | Initiation, branch, and unpaired-base costs for a band-spanning multiloop. |
| $b_C$ | 0.03 | Ordinary-multiloop branch cost for a pseudoknot branch, in addition to PSM. |

The pseudoloop is the region between the crossing bands, including any branches in their gaps. For a pseudoloop, let $m_P$ be the total number of bands, $q_P$ the number of directly attached branches, and $u_P$ the number of unpaired bases directly in its gaps. Each child branch supplies its own internal energy. Density-2 permits chains of three or more bands (Methods). Its internal energy is $E_{P,\text{inside}} = m_P\text{PB} + q_P\text{PPS} + u_P\text{PUP}$; the parent adds PS, PSM, or PSP once according to its loop type.

An ordinary multiloop with $t$ ordinary branches, $k$ pseudoknot branches, and $u$ directly assigned unpaired bases has energy $a_T + b_T(1+t) + c_T u + k(b_C + \mathrm{PSM})$. The loaded Turner initiation, branch, and unpaired-base costs are $a_T = 9.30$, $b_T = -0.90$, and $c_T = 0.00$ kcal/mol; 1 counts the closing pair. For a band-spanning multiloop with $t'$ ordinary side branches, $k'$ pseudoknot side branches, and $u'$ unpaired bases directly in this loop, the energy is $a_p + b_p(2 + t' + k') + c_p u' + k'\mathrm{PSM}$.

The rules below use Boltzmann factors $H$ for a hairpin, $I$ for a stack, bulge, or internal loop, $E$ for an exterior branch, $M$ for an ordinary-multiloop branch, and $C$ for multiloop closure. They include the loaded terminal-pair and neighboring-base terms with dangles=2. The multiloop factors $M$ and $C$ include $b_T$ and $a_T + b_T$, respectively; write $U = B(c_T)$ for an unpaired base. For loops spanning a band, $I^{0.89}$ and $I^{0.74}$ scale stack and internal-loop energies by 0.89 and 0.74, as in CParty. Scaffold-band internal loops have no enumeration-length limit and use the loaded model's long-loop extrapolation.

#### S1.3 Rule summary and correspondence

Table S3 gives parent-to-child rules for all states in Table S1. Multiply each rule's local factor by its children's inside weights. The symbol $u$ denotes a base unpaired in both $G$ and $G'$, and $u^v$ a run of $v \geq 0$ such bases. The symbols $b_<$ and $b_>$ denote the $5'$ and $3'$ bases of one pair. This pair belongs to $G$ or $G'$ in V, to $G'$ in VP, and to $G$ in BE. The run lengths $v_1$ and $v_2$ are also nonnegative. An empty exterior prefix or WI gap has weight 1; WM and WIP require a branch. Right-hand sides show children and bases; VP omits crossed scaffold segments. The symbol □ marks the inner scaffold pair's interior. BE does not count this region; other states supply its weight. Each recursive BE rule shows the outer pair; the BE base rule shows the coincident outer and inner pair. Splits follow Table S1's scaffold regions and Section S1.4's limits. A derivation is a tree of these rules (Section S1.5.1).

Table S3: Structural rules and local Boltzmann factors. Here $u$ is a base unpaired in both $G$ and $G'$, and $u^v$ a run of $v \geq 0$ such bases. The symbols $b_<$ and $b_>$ are the $5'$ and $3'$ bases of a pair: in $G$ or $G'$ for V, in $G'$ for VP, and in $G$ for BE. The symbol □ marks the inner scaffold pair's interior. BE does not count this region; other states supply its weight. Base symbols add no separate factor. Alternatives separated by | use the listed factors in the same order; one factor applies to both alternatives when only one is given. Each child supplies its own interior weight.

| Parent → decomposition | Local factor | Use |
|---|---|---|
| W → W $u$ | 1 | Append one exterior base. |
| W → W V $\mid$ W WMB | $E$ ; $B(\mathrm{PS})$ | Append a branch. |
| W → W WMB W | $B(\mathrm{PS})$ | Root split at a crossed band. |
| V → $b_<\, u^v\, b_>$ | $H$ | Close a hairpin. |
| V → $b_<\, u^{v_1}$ V $u^{v_2}\, b_>$ | $I$ | Stack, bulge, or internal loop. |
| V → $b_<$ VM $b_>$ | 1 | Use a multiloop interior. |
| VM → WM WMv $\mid$ WM WMp | $C$ | At least two branches. |
| VM → $u^v$ WMp | $CU^v$ | One pseudoknot branch. |
| WMv → V; WMp → WMB | $M$ ; $B(b_C + \mathrm{PSM})$ | Start the last branch. |
| $X$ → $X\, u$ | $U$ | $X$: WMv, WMp, or WM. |
| WM → $u^v$ V $\mid$ $u^v$ WMB | $U^v M$ ; $U^v B(b_C + \mathrm{PSM})$ | Start a branch sequence. |
| WM → WM V $\mid$ WM WMB | $M$ ; $B(b_C + \mathrm{PSM})$ | Append a branch. |
| WI → $u$ $\mid$ WI $u$ | $B(\mathrm{PUP})$ | Start or extend an unpaired gap. |
| WI → WI V $\mid$ WI WMB | $B(\mathrm{PPS})$ ; $B(\mathrm{PPS} + \mathrm{PSP})$ | Append a branch. |
| WIP → $u^v$ V $\mid$ $u^v$ WMB | $B(vc_p + b_p)$ ; $B(vc_p + b_p + \mathrm{PSM})$ | First side branch. |
| WIP → WIP V $\mid$ WIP WMB | $B(b_p)$ ; $B(b_p + \mathrm{PSM})$ | Append a side branch. |

| Parent → decomposition | Local factor | Use |
|---|---|---|
| WIP → WIP $u$ | $B(c_p)$ | Append an unpaired base. |
| VPL → $u^v$ VP | $B(vc_p)$ | Left unpaired run, $v \geq 1$. |
| VPR → VP WIP | 1 | Right gap containing a branch. |
| VPR → VP $u^v$ | $B(vc_p)$ | Right unpaired run, $v \geq 1$. |
| VP → $b_<$ WI WI $b_>$ | 1 | Two gaps between crossed bands. |
| VP → $b_<$ WI WI WI $b_>$ | 1 | Three gaps between crossed bands. |
| VP → $b_<$ $u^{v_1}$ VP $u^{v_2}$ $b_>$ | $I^{0.89}$ ; $I^{0.74}$ | Stack or internal loop. |
| VP → $b_<$ WIP VP $b_>$ \| $b_<$ VP WIP $b_>$ | $B(a_p + 2b_p)$ | Branches on one side; other side empty. |
| VP → $b_<$ WIP VPR $b_>$ \| $b_<$ VPL WIP $b_>$ | $B(a_p + 2b_p)$ | Branches on one side; other side nonempty. |
| BE → $b_<$ $\square$ $b_>$ | 1 | Same outer and inner scaffold pair. |
| BE → $b_<$ $u^{v_1}$ BE $u^{v_2}$ $b_>$ | $I^{0.89}$ ; $I^{0.74}$ | Outer scaffold pair; stack or internal loop. |
| BE → $b_<$ WIP BE WIP $b_>$ | $B(a_p + 2b_p)$ | Outer scaffold pair; branches on both sides (Figure S1C). |
| BE → $b_<$ WIP BE $u^v$ $b_>$ \| $b_<$ $u^v$ BE WIP $b_>$ | $B(a_p + 2b_p + vc_p)$ | Outer scaffold pair; branches on one side, other side unpaired. |
| WMBP → VP | $B$(PB) | Start a band chain. |
| WMBP → BE $X$ VP | $B$(2PB) | $X$: WMBP, WMBW, or WI. |
| WMBW → WMBP WI | 1 | Append a nonempty gap. |
| WMB → WMBP | 1 | Require weak closure. |
| WMB → BE WMBP WI | $B$(PB) | Close final band; WI may be empty. |

We regroup CParty's unambiguous density-2 decomposition (Theorem 1 [4]) in a fixed order, preserving split positions and scaffold-pair anchors. Let $\mathrm{BE_{out}}$ be the parent band segment, $\mathrm{BE_{in}}$ its inner continuation, and L and R the left and right segments. The band-spanning multiloop rule becomes

$$\mathrm{BE_{out}} \to b_< \,\mathrm{WIP_L}\,\mathrm{BE_{in}}\,\mathrm{WIP_R}\, b_> \quad \longleftrightarrow \quad \begin{cases} J_\mathrm{R} \to \mathrm{BE_{in}}\,\mathrm{WIP_R} & (1), \\ \mathrm{BE_{out}} \to b_<\,\mathrm{WIP_L}\, J_\mathrm{R}\, b_> & (B(a_p + 2b_p)). \end{cases}$$

For a crossing pair's three WI gaps, numbered from left to right, we use

$$\mathrm{VP} \to b_<\,\mathrm{WI_1}\,\mathrm{WI_2}\,\mathrm{WI_3}\, b_> \quad \longleftrightarrow \quad \begin{cases} J_{23} \to \mathrm{WI_2}\,\mathrm{WI_3} & (1), \\ \mathrm{VP} \to b_<\,\mathrm{WI_1}\, J_{23}\, b_> & (1). \end{cases}$$

Here $J_\mathrm{R}$ and $J_{23}$ are intermediate combinations; parentheses give local factors. Ordinary multiloop closure uses V $\to b_<$ VM $b_>$ followed by VM $\to$ WM WMv (or WMp), placing $C$ only on the latter rule. Shared-band joins retain both anchors; WMB completion requires weak closure. Contraction recovers the original rule and splits; the fixed order gives its unique reverse expansion. Each energy term occurs once, preserving the weight before pruning. Exact calculations evaluate children before parents by interval length; finite beams proceed by right endpoint. Coordinate-level conditions and evaluation procedures are documented in LinearKnot v0.2.2's deduction guide and model description.

#### S1.4 Beam ranking and retention

Inside weight alone can discard partial structures needed for high-weight completed structures. Preliminary inside calculations at width $p = \lceil b^{2/3} \rceil$ estimate the remaining weight needed to reach the root. This estimate is heuristic.

For a state $v$, let $\alpha(v)$ be its inside weight, the sum of its partial-structure weights. A reverse pass gives the log-weight estimate $H_p(v)$ for completing state $v$. For each child, we take the maximum of the parent's estimate plus the local log factor and the other children's inside log weights. The root needs no remaining structure, so $H_p(\mathrm{W}(1, n)) = \log 1 = 0$. For partition-function calculations, the main selection retains the top $b$ candidates of each type at each right endpoint, ranked by $a(v) + H_p(v)$ with $a(v) = \log \alpha(v)$. Structure weights stay unchanged.

Without an estimate, V and exterior WMB starting at $i$ use $a(v) + \log \alpha(\mathrm{W}(1, i-1))$, adding the exterior weight to their left; other types use $a(v)$. Intermediate joins keep the top $b$ for each shared boundary, ranked by child-weight products and available local factors.

Minimum-free-energy prediction replaces summed inside weights with the best partial-structure weights and selects candidates separately. At finite width, these candidates can differ from the partition-function candidates reused by $\gamma$-centroid decoding.

Finite-beam runs use the main-text hairpin-length and outward-search limits $h = d = \max(b, 30)$. The hairpin-length limit does not apply to hairpins fixed in $G$. Exact mode removes $h, d$ and beam selection. Both modes inherit CParty's implemented internal-loop limits to compare the same structure sets: at most $L = 30$ unpaired bases for a new noncrossing V pair and fewer than $L$ for a new crossing VP pair.

Table S4 counts branching levels outside each free base's innermost enclosing scaffold pair. Each level has at least two scaffold branches; the exterior loop counts if it branches, but stacks and internal loops do not. All scaffolds in this table were predicted from LinearPartition pairing probabilities at beam width 200, with the probability threshold 0.7 and minimum helix length two used in Methods. For bpRNA below 400 nt, the stored counts also include the innermost enclosing loop when it branches, so they are upper bounds for the outward-search depth. All maxima are below 30, so $d = 30$ truncates no endpoint search in these datasets.

Table S4: Maximum branching counts in predicted scaffolds over free bases and RNAs, using the counting conventions above.

| Dataset | RNAs | Length (nt) | Maximum levels |
|---|---|---|---|
| bpRNA below 400 nt | 24,625 | 50–399 | 5 |
| bpRNA at least 400 nt | 1,438 | 400–4,381 | 16 |
| RNAcentral long noncoding RNAs | 7 | 4,793–29,106 | 28 |
| NEAT1_1 | 1 | 3,756 | 8 |
| NEAT1_2 | 1 | 22,743 | 26 |

The main run defines probabilities when its root weight is positive.

#### S1.5 Probability outputs

##### S1.5.1 Inside and outside after beam pruning

For state $v$, let $\mathcal{R}(v)$ be the set of allowed rules whose parent and children survive beam pruning. Rule $r$ has local Boltzmann factor $\omega_r$ and $k_r$ children $u_{r,1}, \ldots, u_{r,k_r}$; $t$ and $s$ index these children. The inside weights satisfy

$$\alpha(v) = \sum_{r \in \mathcal{R}(v)} \omega_r \prod_{s=1}^{k_r} \alpha(u_{r,s}).$$

An empty product is 1, and the root $v_0 = \mathrm{W}(1, n)$ gives $\hat{Z} = \alpha(v_0)$. Finite-beam calculations reuse inside selections; exact calculations use all allowed rules. Sums are evaluated in log space.

The outside weight $\beta(v)$ sums weights surrounding $v$ and connecting it to the root. Initialize $\beta(v_0) = 1$ and other outside weights to 0. In reverse inside order, each rule with parent $v$ contributes

$$\beta(u_{r,t}) \mathrel{+}= \beta(v)\omega_r \prod_{\substack{s=1 \\ s \neq t}}^{k_r} \alpha(u_{r,s}), \qquad 1 \leq t \leq k_r.$$

The outside weight $\beta(v)$ is also the derivative $\partial \hat{Z} / \partial \alpha(v)$ [3]. Figure S2 illustrates the two directions. The probability of using rule $r$ is

$$\pi_r = \frac{\beta(v)\omega_r \prod_{s=1}^{k_r} \alpha(u_{r,s})}{\hat{Z}} = \frac{\partial \log \hat{Z}}{\partial \log \omega_r}.$$

Summing $\pi_r$ over rules that introduce an event once per derivation gives its probability. Exact mode replaces $\hat{Z}$ by $Z$.

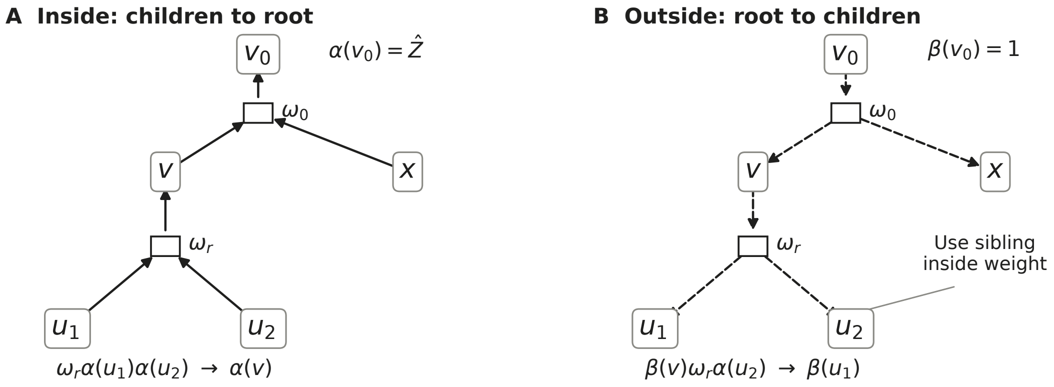


Figure S2: Inside and outside flow through a binary decomposition. Each square joins a parent and two children with local factor $\omega$. A: rule $r$ adds $\omega_r\alpha(u_1)\alpha(u_2)$ to the parent's inside weight $\alpha(v)$. B: rule $r$ adds $\beta(v)\omega_r\alpha(u_2)$ to the child's outside weight $\beta(u_1)$. Contributions are summed over retained rules. This schematic follows Eisner's inside/outside interpretation [3].

##### S1.5.2 Pair and context probabilities

Each new pair is introduced once at a V or VP parent; summing its introducing rules' $\pi_r$ gives $p_{ij} = p_{ji}$. The unpaired probability $q_i$ follows Methods. Of the seven contexts in Methods, PseudoLoop covers gaps between crossing bands (cost PUP); unpaired segments of band-spanning multiloops are Multiloop (cost $c_p$). Hairpin, Bulge, and Internal retain their types inside pseudoknots.

Each rule assigns its added unpaired bases once to its loop type: enclosed hairpin, bulge, or internal loop; Exterior for W; PseudoLoop for WI; Multiloop for multiloop extensions. Summing $\pi_r$ for assignments of base $i$ to context $c$ gives $P_i(c)$, using cumulative sums for intervals. Paired bases are Stem. Enumeration verifies these assignments, $P_i(\text{Stem}) = \sum_{j\neq i} p_{ij}$, and $\sum_{c\neq\text{Stem}} P_i(c) = q_i$.

###### S1.5.3 Total accessibility and accessibility by loop type

For window $W_{s,w} = [s, s+w-1]$, $1 \le s \le n-w+1$, let $C_R(i)$ be the context at base $i$ in structure $R$. In addition to total accessibility $A(s,w)$, define $A_c(s,w) = \widehat{\Pr}\{W_{s,w}$ is unpaired and $C_R(i) = c$ throughout the window$\}$ for each of the six unpaired contexts. A contiguous unpaired window has one context, including across separately assigned PseudoLoop segments. Thus $\sum_{c\neq\text{Stem}} A_c(s,w) = A(s,w)$ and $A_c(s,1) = P_s(c)$.

For a rule assigning an entire unpaired interval $[a,z]$, add $\pi_r$ to window starts $a \le s \le z-w+1$. For a window spanning successive one-base extensions of state type $X$ at fixed left boundary $i$, let $g_t$ be the Boltzmann factor for extending through unpaired base $t$. Let $\alpha_{\mathrm{u}}(X(i,s))$ be the sum of inside contributions from steps that enter $X(i,s)$ with base $s$ unpaired. With $e = s+w-1$, the chain contributes

$$\alpha_{\mathrm{u}}(X(i,s)) \left(\prod_{t=s+1}^{e} g_t\right) \frac{\beta(X(i,e))}{\hat{Z}}.$$

###### S1.5.4 Error bound from the retained weight

For fixed $S, G$, assume $\widehat{\Omega} \subseteq \Omega$, one count per structure, unchanged weights, and positive $Z, \hat{Z}$. Methods derives $\mathrm{TV}(\widehat{\Pr}, \Pr) = 1-\rho$, with $\rho = \hat{Z}/Z = \exp(-\Delta F/\kappa)$ and $\Delta F = \hat{F} - F$. Here $F = -\kappa \log Z$ and $\hat{F} = -\kappa \log \hat{Z}$ are the exact and finite-beam ensemble free energies. To obtain a marginal distribution, partition structures into disjoint sets $A_k$, such as the pairing partners and unpaired state of one base. Writing $d(R) = \widehat{\Pr}(R) - \Pr(R)$, the triangle inequality gives

$$\frac{1}{2}\sum_k \left|\sum_{R\in A_k} d(R)\right| \le \frac{1}{2}\sum_k \sum_{R\in A_k} |d(R)| = \frac{1}{2}\sum_R |d(R)| = 1-\rho.$$

This bounds pairing and context TV at each base, and their RNA-level averages. Let $\mathcal{A} \subseteq \Omega$ be an event, such as the set of structures with an entirely unpaired window. Both distributions sum to 1, so $\sum_{R\in\Omega} d(R) = 0$ and

$$\sum_{R\in\mathcal{A}} d(R) = -\sum_{R\in\Omega\setminus\mathcal{A}} d(R).$$

The two sums have equal absolute values, giving

$$\begin{aligned}
|\widehat{\Pr}(\mathcal{A}) - \Pr(\mathcal{A})| &= \frac{1}{2}\left[\left|\sum_{R\in\mathcal{A}} d(R)\right| + \left|\sum_{R\in\Omega\setminus\mathcal{A}} d(R)\right|\right] \\
&\le \frac{1}{2}\left[\sum_{R\in\mathcal{A}} |d(R)| + \sum_{R\in\Omega\setminus\mathcal{A}} |d(R)|\right] = \frac{1}{2}\sum_{R\in\Omega} |d(R)| = 1-\rho.
\end{aligned}$$

Thus the absolute accessibility error is at most $1-\rho$.

Within each $A_k$, positive and negative differences can cancel; equality requires one sign per set. For 300–399 nt at $b=200$, an RNA with $\Delta F = 0.0111$ kcal/mol (the RNA-level 90th percentile, or p90) has bound about 0.018, whereas the BPP mean-TV p90 is 0.0015 (Table S5). The bound requires exact $Z$, excluding the $b=400$ reference, and does not bound relative or opening-energy errors.

#### S1.6 Sampling and $\gamma$-centroid

Sampling starts at $\mathrm{W}(1,n)$ and chooses each rule in proportion to its local factor times its children's inside weights.

Finite-beam $\gamma$-centroid decoding uses retained partition-function candidates without further pruning. For candidate $Y$ and ensemble structure $R$, let $m(Y,R)$ be the number of pairs in both and $u(Y,R)$ the number of pairs in neither, over all $1 \le i < j \le n$. Here $p_{ij}$ is the base-pairing probability. The $\gamma$-centroid estimator maximizes [6]

$$\mathbb{E}_R[\gamma m(Y,R) + u(Y,R)] = C + \sum_{(i,j)\in Y} \big[(\gamma+1)p_{ij} - 1\big], \qquad C = \sum_{i<j}(1-p_{ij}).$$

Since $C$ is independent of $Y$, this is the pair-score objective in Methods; scaffold pairs also add the same constant to every candidate. With $\gamma = 1$, the estimator minimizes the expected number of missed and extra pairs.

#### S1.7 User interface

From the LinearKnot v0.2.2 source directory, `pip install .` installs the `linearknot` command. Each input record has an optional `>name` line, a sequence, and a scaffold of the same length, using balanced round brackets and dots:

```
>example
GGGGGAACCAACCC
((.....)).....
```

Save records in `input.txt` and run `linearknot fold -i input.txt --bpp bpp.tsv`. Files may contain multiple records; extra bracket types mark output crossings.

| Command | Output |
|---|---|
| `fold` | Minimum-energy structure and ensemble free energy; `--bpp FILE` saves BPPs. |
| `profile` | Seven structural-context probabilities per base. |
| `access` | Window opening energies; `--probability` gives joint unpaired probabilities and `--by-loop` adds loop contributions. |
| `sample` | Boltzmann samples; `--count K` sets the number of draws. |
| `subopt` | Up to `--top-k K` distinct structures in energy order. |
| `mea` | One maximum expected accuracy structure. |
| `centroid` | One $\gamma$-centroid structure. |

All commands accept `-i input.txt` and `--exact` (no beam pruning or beam-only limits). Defaults are $b = 100$ and $h = d = 30$; for the paper's $b = h = d = 100$, add `--beam 100 --max-hairpin 100 --max-case3-depth 100`.

The scaffold helper uses the paper's rule by default: LinearPartition-V, beam 200, dangles=2, pair probability $\geq 0.7$, and at least two stacked pairs per helix. After preparing LinearPartition as described in the README, run

```
python3 scripts/scaffold/make_scaffold.py sequences.fa > input.txt
```

From Python, call the `linearknot` command through `subprocess`. This example iterates over sample output lines, including headers:

```
import subprocess
cmd = ["linearknot", "sample", "-i", "input.txt",
       "--count", "20", "--seed", "42"]
with subprocess.Popen(cmd, stdout=subprocess.PIPE, text=True) as proc:
    for line in proc.stdout:
        print(line, end="")
    if proc.wait():
        raise RuntimeError("linearknot failed")
```

For suboptimal output, set `cmd` to

```
["linearknot", "subopt", "-i", "input.txt", "--top-k", "8"]
```

Both commands write one structure at a time, retaining graph memory (Section S2.3). `sample --unique` instead reports counts by structure.

To reproduce the computational experiments, see the LinearKnotExperiments README. The CParty reference uses source `800e95a` with corrections in the experiment manifest. Our usual ensemble requires completed-region weak closure; comparisons omit it in both programs to match CParty, with Turner 2004, 37 °C, dangles=2, exact mode, and matched internal-loop limits.

### S2 Computational cost

Let $n$ be sequence length, $b$ beam width, and $r_w$ the number of requested accessibility window lengths. Costs exclude scaffold prediction; space is maximum simultaneous storage. We assume constant-time arithmetic on fixed-size numbers and expected constant-time hash lookups, which search for stored states by their keys. Expected time refers to these lookups, rather than an average over RNA inputs.

#### S2.1 Finite-beam probability calculations and decoding

**Proposition (finite-beam cost).** Assume $2 \leq b \leq n$, fixed local limits $L, h, d$, width $b$ for the state and intermediate selections, and the standard procedure for retaining a derivation of $G$ alone (Section S1.4). For any supplied pseudoknot-free scaffold, the partition function, pairing and seven context probabilities, and finite-beam MFE or $\gamma$-centroid decoding each take expected time $T_0 = O(nb^2 \log b)$ and space $S_0 = O(nb)$, including ranking, retention of $G$ alone, rule reconstruction, and output. Constants may depend on $L, h, d$; increasing these limits with $b$ can increase them.

At each endpoint, bounded local searches and joins of retained lists generate $O(b^2)$ candidate combinations, whose selection costs $O(b^2 \log b)$. Over $n$ endpoints this gives $T_0$, which also covers outside rule regeneration and pairing and context aggregation.

Let $c_{\max} \leq n$ be the maximum number of children in the scaffold nesting tree, and $E_d \leq nd$ the total outward branching levels stored over free endpoints. Retained states and temporary joins use $O(nb + E_d)$ and $O(b^2 + c_{\max}b)$ space, respectively. Their sum is $O(nb)$ for fixed $d$ and $b \leq n$. The preliminary ranking also fits these bounds, since $p = \lceil b^{2/3} \rceil \leq b$.

Accessibility adds window aggregation after outside propagation. For total accessibility or accessibility by loop type, each requested window length adds $O(nb^2)$ work and $O(n)$ output space. The expected time $T_A$ and space $S_A$ are

$$T_A = O\big(nb^2 \log b + r_w nb^2\big) , \qquad S_A = O(nb + nr_w).$$

Six unpaired loop types change only a constant factor. These bounds are linear in $n$ for fixed $b, L, h, d, r_w$, as in our experiments with $L = 30$ and $h = d = \max(b, 30)$ at each chosen $b$.

##### S2.2 Exact numerical probabilities

Let $S_{\text{ex}} = O(n^2)$ be the number of exact states, including BE scaffold-pair combinations, and $D_{\text{ex}}$ the number of generated rule applications. Let $T_{\text{ex}}$ be the full inside cost, including scaffold geometry and rule generation. Pairing and context probabilities regenerate rules for outside and aggregation, taking expected time $O(T_{\text{ex}} + D_{\text{ex}} + S_{\text{ex}} + n)$ and space $O(n^2)$ at fixed local-loop limits. The finite-beam bound does not give a cubic bound for exact scaffold searches in $T_{\text{ex}}$.

With accessibility aggregation for each window length, expected time is $O(T_{\text{ex}} + (r_w + 1)D_{\text{ex}} + r_w S_{\text{ex}} + nr_w)$ and space is $O(n^2 + nr_w)$. These bounds describe numerical probabilities; they do not cover the additional traceback storage used by exact MFE or exact centroid decoding.

##### S2.3 Stored graphs and structure output

Suboptimal enumeration lists structures in order of increasing energy. Finite-beam sampling and suboptimal enumeration save rule graphs; exact sampling saves the operations used to sum weights. This storage is outside the numerical $O(nb)$ bound, and sampling may scan several rules per state.

Writing $k_s$ structures of length $n$ takes $\Omega(k_s n)$ time and $O(n)$ completed-output space when delivered one at a time, or $O(k_s n)$ output space when collected. Saved graphs remain; suboptimal searches may store more information as outputs increase, and sample-frequency aggregation stores distinct structures and counts.

#### S3 Supplementary results

##### S3.1 Validation details

All 305 inputs (300 bpRNA sequences of 52–396 nt and five boundary and branching tests) agreed with CParty within $10^{-9}$ in $\log Z$ and $10^{-10}$ kcal/mol in free energy under Section S1.7's settings. The maximum absolute differences were $1.9 \times 10^{-12}$ in $\log Z$ and $1.194 \times 10^{-12}$ kcal/mol in ensemble free energy.

With an empty scaffold and matched parameters, ViennaRNA 2.7.2 [8] agreed on 337 sequences within $1.450 \times 10^{-12}$ in $\log Z$ and $1.246 \times 10^{-12}$ in BPPs. Single-structure energies (548 conditions) and independent MFE rescoring agreed within $2.843 \times 10^{-14}$ kcal/mol.

Enumeration of 156,534 structures across 729 cases on 706 inputs (5–20 nt) gave one derivation per structure and unique contexts. Partition functions agreed with complete enumeration using the same rules and local weights within $2.14 \times 10^{-14}$ in $\log Z$. BPPs, context probabilities, and total accessibility agreed with enumeration within $2.2 \times 10^{-14}$, MFE energies within $3.6 \times 10^{-15}$ kcal/mol, and $\gamma$-centroid scores within $1.8 \times 10^{-13}$ in 6,561 comparisons using their respective retained ensembles.

Sampling uses six inputs in exact and $b = 1, 3$ modes, with $N =$ 200,000 draws per condition (3,600,000 total). For $M = 972$ structure-frequency events and allowed simultaneous error probability $\alpha_{\text{err}} = 0.01$, the simultaneous frequency-error bound is $\epsilon = \sqrt{\frac{\log \frac{2M}{\alpha_{\text{err}}}}{2N}} = 0.005518$. Hoeffding's inequality [7] and a sum over events give this bound without event independence. Sampling frequencies differed from the exact probabilities of their respective retained ensembles by at most 0.0030, below the simultaneous bound 0.005518. The maximum TV between sampled and theoretical distributions was 0.00439478.

#### S3.2 Approximation accuracy

Comparisons of total accessibility vary $h = d = \max(b, 30)$ with $b$.

Table S5: RNA-level p90 and maximum differences, with exact references below 400 nt and finite $b = 400$ above. $\Delta F$ is signed (kcal/mol); BPP and context columns give mean TV over free bases and accessibility gives mean absolute difference over eligible windows. RNA counts are shown separately for all sequences and those with eligible 16-nt windows.

| Length (nt) | RNAs | RNAs with 16-nt windows | $b$ | Statistic | $\Delta F$ | BPP TV | Context TV | 16-nt accessibility |
|---|---|---|---|---|---|---|---|---|
| Exact reference, length below 400 nt | | | | | | | | |
| 50–99 | 12,428 | 5,937 | 200 | p90 | $1.07 \times 10^{-14}$ | $1.08 \times 10^{-14}$ | $1.92 \times 10^{-14}$ | $1.4 \times 10^{-15}$ |
| | | | | Maximum | $9.59 \times 10^{-14}$ | $1.79 \times 10^{-13}$ | $1.87 \times 10^{-13}$ | $5.57 \times 10^{-14}$ |
| 100–199 | 9,656 | 5,862 | 200 | p90 | $2.84 \times 10^{-14}$ | $3.81 \times 10^{-14}$ | $6.06 \times 10^{-14}$ | $4.67 \times 10^{-15}$ |
| | | | | Maximum | $3.27 \times 10^{-13}$ | $4.9 \times 10^{-13}$ | $3.62 \times 10^{-13}$ | $1.71 \times 10^{-13}$ |
| 200–299 | 1,446 | 1,213 | 200 | p90 | $8.53 \times 10^{-14}$ | $1.83 \times 10^{-13}$ | $2.07 \times 10^{-13}$ | $8.45 \times 10^{-15}$ |
| | | | | Maximum | 0.3 | 0.0279 | 0.0314 | 0.000973 |
| 300–399 | 1,095 | 1,048 | 200 | p90 | 0.0111 | 0.0015 | 0.00174 | $2.48 \times 10^{-5}$ |
| | | | | Maximum | 3.89 | 0.373 | 0.206 | 0.0305 |
| Finite reference ($b = 400$), length at least 400 nt | | | | | | | | |
| 400–799 | 584 | 572 | 300 | p90 | 0.0167 | 0.0019 | 0.00194 | $2.93 \times 10^{-5}$ |
| | | | | Maximum | 3.87 | 0.416 | 0.275 | 0.0249 |
| 800–1599 | 423 | 423 | 300 | p90 | 1.02 | 0.0832 | 0.0648 | 0.00102 |
| | | | | Maximum | 6.39 | 0.347 | 0.227 | 0.0195 |
| 1600–3199 | 340 | 340 | 300 | p90 | 2.68 | 0.104 | 0.0659 | 0.000986 |
| | | | | Maximum | 13 | 0.35 | 0.24 | 0.0479 |
| 3200–4381 | 91 | 91 | 300 | p90 | 3.35 | 0.0791 | 0.0548 | 0.00318 |
| | | | | Maximum | 8.72 | 0.154 | 0.0909 | 0.00858 |

Pruning with unchanged weights gives $\hat{Z} \leq Z$ and $\hat{F} \geq F$ before rounding (Section S1.5.4). Larger negative differences against $b = 400$ are possible because its retained set need not include all structures retained at smaller widths.

The sum of exact probabilities of pairs crossing $G$ is their expected count. We grouped RNAs of 50–399 nt by this count: below 0.1, from 0.1 to below 1, or at least 1. The last group had larger 90th percentiles of pairing and context TV averaged over free bases within each RNA at $b = 20, 50, 100$. At $b = 200$, these percentiles were below $7 \times 10^{-14}$ in all three groups. Each percentile pools lengths within its group; the groups differed in length composition.

Table S6 counts large local decreases and increases without selecting coordinates by exact probability. All coordinates are included, with missing pairs assigned zero; adjacent events are dependent. Neither change occurred in 23,530 shorter RNAs at $b = 200$.

Table S6: Large local changes in 1,095 RNAs of 300–399 nt. Decrease: exact probability $\geq 0.5$ and beam probability $< 0.1$; increase: exact $< 0.1$ and beam $\geq 0.5$. Rows measure pair probability, single-base PseudoLoop probability, or total four-base accessibility. Counts give events and affected RNAs; percentages use 1,095.

| | | Decrease | | Increase | |
|---|---|---|---|---|---|
| Event | $b$ | Events | RNAs (%) | Events | RNAs (%) |
| All pairs | 100 | 1,095 | 79 (7.21%) | 404 | 38 (3.47%) |
| | 200 | 117 | 10 (0.91%) | 8 | 5 (0.46%) |
| Pairs crossing $G$ | 100 | 129 | 26 (2.37%) | 16 | 5 (0.46%) |
| | 200 | 27 | 7 (0.64%) | 2 | 2 (0.18%) |
| PseudoLoop bases | 100 | 306 | 42 (3.84%) | 14 | 4 (0.37%) |
| | 200 | 80 | 15 (1.37%) | 0 | 0 (0.00%) |
| 4-nt windows | 100 | 118 | 43 (3.93%) | 154 | 41 (3.74%) |
| | 200 | 10 | 7 (0.64%) | 8 | 4 (0.37%) |

We did not analyze opening-energy differences.

##### S3.3 Time and memory

All 63 real-RNA and 1,404 synthetic measurements completed on seven RNAcentral sequence–scaffold inputs and 156 synthetic controls. Hardware and repetitions follow Methods, with an 8-GiB per-process virtual-memory limit and randomized order. Release builds used the GNU Compiler Collection 11.5.0; timing covers startup through output.

Table S7: RNAcentral whole-process resource medians at $b = h = d = 100$. The three calculations give partition function, BPPs, and MFE; loop profiles; and total accessibility with loop profiles. All seven are annotated long noncoding RNAs (annotations retrieved 8 October 2026).

| | | Wall time (s) | | | Peak memory (GiB) | | |
|---|---|---|---|---|---|---|---|
| RNAcentral identifier | nt | BPP, MFE | Profiles | Accessibility | BPP, MFE | Profiles | Accessibility |
| URS000233CDDE | 4,793 | 60.21 | 39.15 | 39.28 | 0.59 | 0.55 | 0.55 |
| URS00019626CE | 5,482 | 112.54 | 75.95 | 82.39 | 0.88 | 0.86 | 0.86 |
| URS000239D8A4 | 6,990 | 126.08 | 86.75 | 82.45 | 0.97 | 0.92 | 0.93 |
| URS00023C9544 | 11,832 | 159.69 | 104.19 | 103.84 | 1.42 | 1.35 | 1.35 |
| URS000241043C | 13,652 | 301.54 | 211.73 | 204.71 | 2.24 | 2.15 | 2.15 |
| URS000238C72C | 22,543 | 423.97 | 279.78 | 280.84 | 3.05 | 2.94 | 2.94 |
| URS00023A4DB9 | 29,106 | 413.95 | 275.78 | 284.40 | 3.22 | 3.04 | 3.04 |

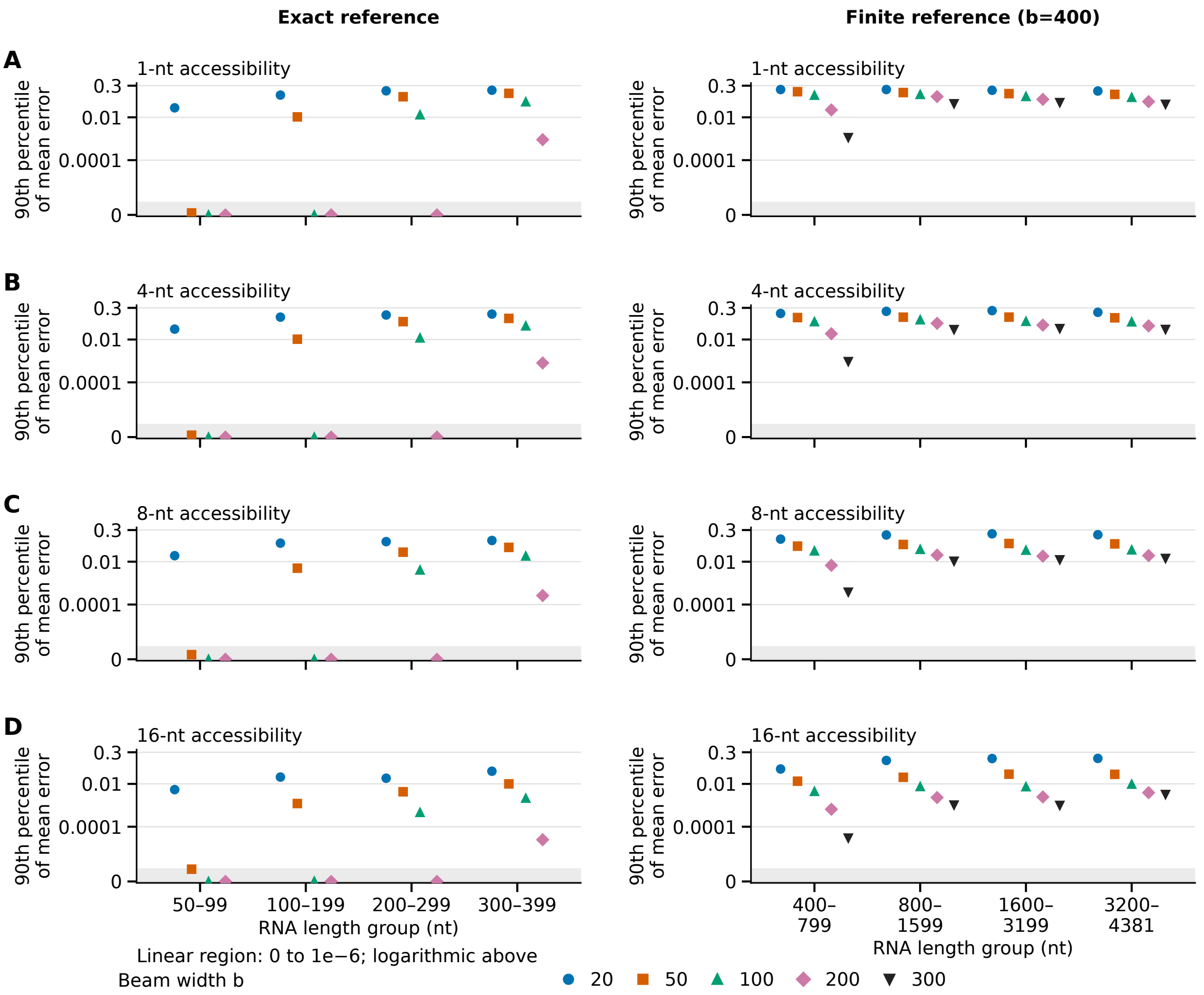


Figure S3: Total-accessibility errors: bpRNA below 400 nt against exact calculations (left); 400–4,381 nt against $b = 400$ (right). A–D use 1-, 4-, 8-, and 16-nt windows, with $h = d = \max(b, 30)$. For each RNA, we average absolute probability differences over windows with no base paired in $G$, including zeros. Points show the 90th percentile of these averages across RNAs. RNA counts: 24,625, 24,592, 22,322, and 14,060 (left); 1,438 for 1–8 nt, 1,426 for 16 nt (right). Shading is linear (0–0.000001); higher values are logarithmic. Symbols follow Figure 2.

On 40-core allocations on different hosts, concurrency controls compared one and up to nine processes for three output groups on 4,793-, 5,482-, and 6,990-nt inputs, using medians of three measurements after one warm-up. Across the nine input–output pairs, concurrent to single-process time ratios were 1.01–1.26 (median 1.04) and peak memory ratios 0.999–1.005; we did not correct the main results for these ratios.

The synthetic set has seven patterns: empty scaffold, one long stem, three separated helices, three helices with free bases in one gap, 12 separated helices, nested multiloops, and balanced multiloops. Nonempty scaffolds have $\lfloor 0.15n \rfloor$ pairs; the two multiloop shapes match pair and branch counts, with binary branching. Lengths 250, 500, 1,000, 2,000, and 4,000 nt, with three independently seeded sequences per shape and length, give 105 inputs (Figure S4). Scaffold pairs use AU, UA,

GC, or CG, with 50% GC overall. Another 51 inputs probe branching levels near $d = 30, 100$ and are not shown in the figure. The run configuration specifies inputs and three measurement repetitions; generation notes record seeds, gap allocation, and branching levels.

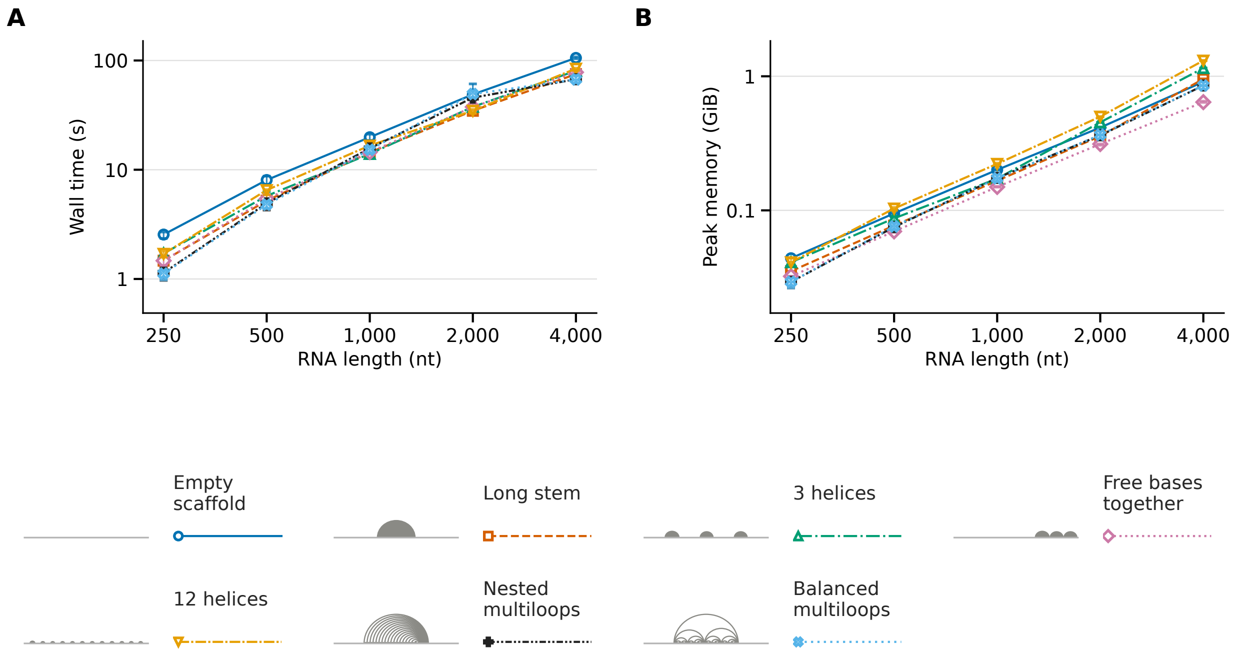


Figure S4: Resource costs for seven synthetic scaffold shapes. A: wall time; B: peak memory for partition function, BPPs, and MFE. Axes use logarithms (base 2 for length, base 10 for time and peak memory). Symbols are medians of three sequence medians, each from three measurements; bars span all nine and may be smaller than symbols.

From 2,000 to 4,000 nt, the costs in Figure S4A and B increased by factors of 1.36–2.43 for time and 2.05–2.67 for peak memory.

##### S3.4 Full-length NEAT1 calculations

NEAT1_1 (NR_028272.1, 3,756 nt) has 727 scaffold pairs; NEAT1_2 (NR_131012.1, 22,743 nt) has 4,161. Both use the scaffold construction rule in the main-text Methods.

##### S3.5 Reference-pair recovery in exact calculations

Recovery means predicting a pair present in the reference annotation. We randomly selected 200 of 982 bpRNA-1m(90) inputs of 50–300 nt with at least two reference pairs recorded as `[` and `]` in the bpRNA annotation [2], using scaffolds from the nested reference pairs. In exact calculations, we compared the existing Turner 2004 [9] and DP09 [1] parameter sets for ordinary loops, keeping CParty's pseudoknot constants fixed (Table S9).

Table S8: NEAT1 whole-process resources at $b = h = d = 100$. Each cell is one measurement. Loop-type accessibility includes contexts and total accessibility and its six components at lengths 1, 4, 8, and 16; other outputs follow Table S7. Centroid uses $\gamma = 1$.

| | NEAT1_1 | | NEAT1_2 | |
|---|---|---|---|---|
| Analysis | Time (s) | Peak memory (GiB) | Time (s) | Peak memory (GiB) |
| BPP and MFE | 53 | 0.49 | 392 | 2.96 |
| Loop profiles | 35 | 0.47 | 245 | 2.79 |
| Total accessibility | 40 | 0.47 | 255 | 2.79 |
| Loop-type accessibility | 36 | 0.47 | 247 | 2.80 |
| 1,000 samples | 53 | 1.96 | 356 | 15.47 |
| $\gamma$-centroid | 75 | 0.50 | 526 | 3.03 |

Table S9: Reference-pair recovery with exact calculations on 200 RNAs. Ordinary loops (hairpins, stacks, bulges, internal loops, and multiloops) use the published Turner 2004 or DP09 parameter set; CParty's pseudoknot constants are the same in both calculations. We did not tune or fit any parameter for this comparison. Targets are canonical reference pairs crossing $G$; matches join the same two positions in the predicted MFE structure. Sensitivity divides matches by reference targets; positive predictive value divides matches by predicted pairs crossing $G$, summing counts across RNAs. Mean pair probabilities use these targets within each RNA.

| Measure | Turner 2004 ordinary loops | DP09 ordinary loops |
|---|---|---|
| Pairs with probability $> 0.5$ | 179 of 1,040 (17.2%) | 80 of 1,040 (7.7%) |
| RNAs with mean pair probability $> 0.5$ | 34 of 200 | 13 of 200 |
| Sensitivity | 210 of 1,040 (20.19%) | 110 of 1,040 (10.58%) |
| Positive predictive value | 210 of 850 (24.71%) | 110 of 413 (26.63%) |

#### S3.6 Resource comparisons with CParty and Spark

Table S10 compares partition functions on 105 synthetic inputs and 107 real RNAs under Section S1.7's settings, with LinearKnot exact or $b = h = d = 100$. Timing covers startup through postprocessing, excluding sampling, MFE, and outside calculations. CParty was faster through 1,000 nt and LinearKnot $b = 100$ at 2,000 nt on common synthetic successes. All 51 common CParty and exact LinearKnot successes met Section S3.1's tolerances; synthetic exact was measured through 1,000 nt.

The real RNA set has 20 bpRNA sequences per group (seed 20261025), within ±10% of 250, 500, 1,000, 2,000, and 4,000 nt, plus the seven RNAcentral inputs, all with saved scaffolds. Selected ranges were 225–275, 451–550, 905–1,077, 1,805–2,158, and 3,607–4,065 nt. We compiled LinearKnot v0.2.1, CParty, and Spark `3105c311` with the same compiler in release mode. Each input and method had one warm-up and three measurements in randomized order. On 16-core allocations, up to eight single-threaded processes ran concurrently through 2,158 nt (2,000 nt for synthetic inputs) and four above, except for the long RNAcentral runs described below. CParty and Spark ran alone when their memory limit was 32 GiB, for RNAcentral inputs of 11,832–29,106 nt. Methods could use different hosts above 1,077 nt.

The five bpRNA groups had virtual-memory limits of 1, 1, 4, 4, and 8 GiB, and CParty and Spark time limits of 10, 30, 120, 180, and 600 s. LinearKnot $b = 100$ had 300 s in the last group; exact was measured through 2,158 nt, with 900 s in the 2,000-nt group. For RNAcentral, CParty and Spark had 900 s and 8 GiB through 6,990 nt, then 3,600 s and 32 GiB from 11,832 nt; LinearKnot $b = 100$ had 900 s and 8 GiB throughout, with or without MFE.

Partition-function medians require three successful runs with finite partition-function values. Completion denominators include failures. All 177 finite CParty and exact LinearKnot measurement pairs (59 real RNAs) passed the partition-function tolerances: maximum $|\Delta \log Z| = 1.18 \times 10^{-11}$ and energy difference $7.30 \times 10^{-12}$ kcal/mol.

Table S10: Partition-function time (s), peak memory (MiB), and completed inputs for CParty and LinearKnot. Values are medians of input medians: synthetic resources use common successes, real-RNA resources each method's successful set. At 4,000 nt, synthetic resources use only LinearKnot $b = 100$. Each length group contains 21 synthetic inputs or 20 bpRNA inputs; RNAcentral has one input at each length. Dashes mean no finite result or unmeasured exact mode.

| | CParty | | | LinearKnot exact | | | LinearKnot $b = 100$ | | |
|---|---|---|---|---|---|---|---|---|---|
| Group (nt) | s | MiB | Done | s | MiB | Done | s | MiB | Done |
| Synthetic | | | | | | | | | |
| 250 | 0.067 | 9.0 | 21 | 0.367 | 16.0 | 21 | 0.818 | 30.0 | 21 |
| 500 | 0.367 | 24.0 | 21 | 2.67 | 53.2 | 21 | 2.82 | 62.1 | 21 |
| 1,000 | 4.13 | 84.2 | 9 | 15.95 | 201.3 | 21 | 8.28 | 139.8 | 21 |
| 2,000 | 34.97 | 330.6 | 6 | – | – | – | 23.40 | 298.5 | 21 |
| 4,000 | – | – | 0 | – | – | – | 47.58 | 931.1 | 21 |
| Real RNA: bpRNA | | | | | | | | | |
| 250 | 0.116 | 9 | 20 | 0.217 | 16.6 | 20 | 0.417 | 21 | 20 |
| 500 | 0.367 | 24 | 20 | 1.19 | 53.2 | 20 | 2.35 | 61.5 | 20 |
| 1,000 | 2.52 | 78 | 19 | 10.5 | 186 | 20 | 7.38 | 140 | 20 |
| 2,000 | – | – | 0 | 52.8 | 743 | 20 | 12.8 | 216 | 20 |
| 4,000 | – | – | 0 | – | – | – | 26.7 | 444 | 20 |

RNAcentral: LinearKnot $b = 100$ completed all seven; CParty completed none; exact was not measured.

| Length (nt) | 4,793 | 5,482 | 6,990 | 11,832 | 13,652 | 22,543 | 29,106 |
|---|---|---|---|---|---|---|---|
| Time (s) | 32.2 | 55.7 | 65.8 | 87.2 | 157 | 240 | 219 |
| Memory (MiB) | 549 | 751 | 939 | 1,310 | 1,940 | 2,860 | 3,190 |

All synthetic CParty failures were non-finite results in all four runs. The 48 failures among 107 real RNAs were 43 non-finite results (994 nt, all 40 inputs at 1,805–4,065 nt, and 4,793 and 5,482 nt), one time limit (6,990 nt), three with time limits or crashes across repetitions (11,832, 13,652, 22,543 nt), and one 32-GiB memory limit (29,106 nt). Measurements gave 129 non-finite results, seven time limits, five crashes of unknown cause, and three memory limits. Non-finite results reflect numerical range, not an algorithmic limit.

CParty's partition function was measured through a program that reads its inside result, with sampling disabled and scaling seed set to zero. Scaling multiplies intermediate values to avoid numerical overflow. In a preliminary check, enabling it gave different recovered partition functions after undoing the scaling (experiment record). We therefore used no scaling, and long inputs overflowed. Correcting the scaling could avoid these overflows, but we did not measure a corrected version.

Spark sparsifies HFold's recurrences to compute MFE [5], while LinearKnot computes the partition function and MFE at $b = h = d = 100$. Spark's resource use divided by LinearKnot's was 0.0372–

0.424 for time and 0.0244–0.286 for peak memory across bpRNA group medians, and 0.562–4.62 for time and 0.0284–0.0626 for memory on completed RNAcentral inputs (medians of paired input ratios). Spark completed 99 of 100 bpRNA inputs (19 of 20 in the 500-nt group and all inputs in other groups) and the four RNAcentral inputs through 11,832 nt; LinearKnot completed all 107 inputs.

#### S3.7 Sensitivity to beam width and local limits on long RNAs

We varied $b = 25, 50, 75, 100, 200, 300, 400$ with $h = d = \max(b, 30)$ on the seven RNAcentral inputs in Table S7 (Figure S5). All 49 settings completed with LinearKnot v0.2.1. This measures sensitivity to beam width and local limits together. We plotted ensemble free energy per base, $-R_{\text{gas}}T \log \hat{Z}_b/n$, where $\hat{Z}_b$ is the approximate partition function at width $b$ and $n$ is sequence length. The gas constant $R_{\text{gas}}$ and absolute temperature $T$ follow Methods.

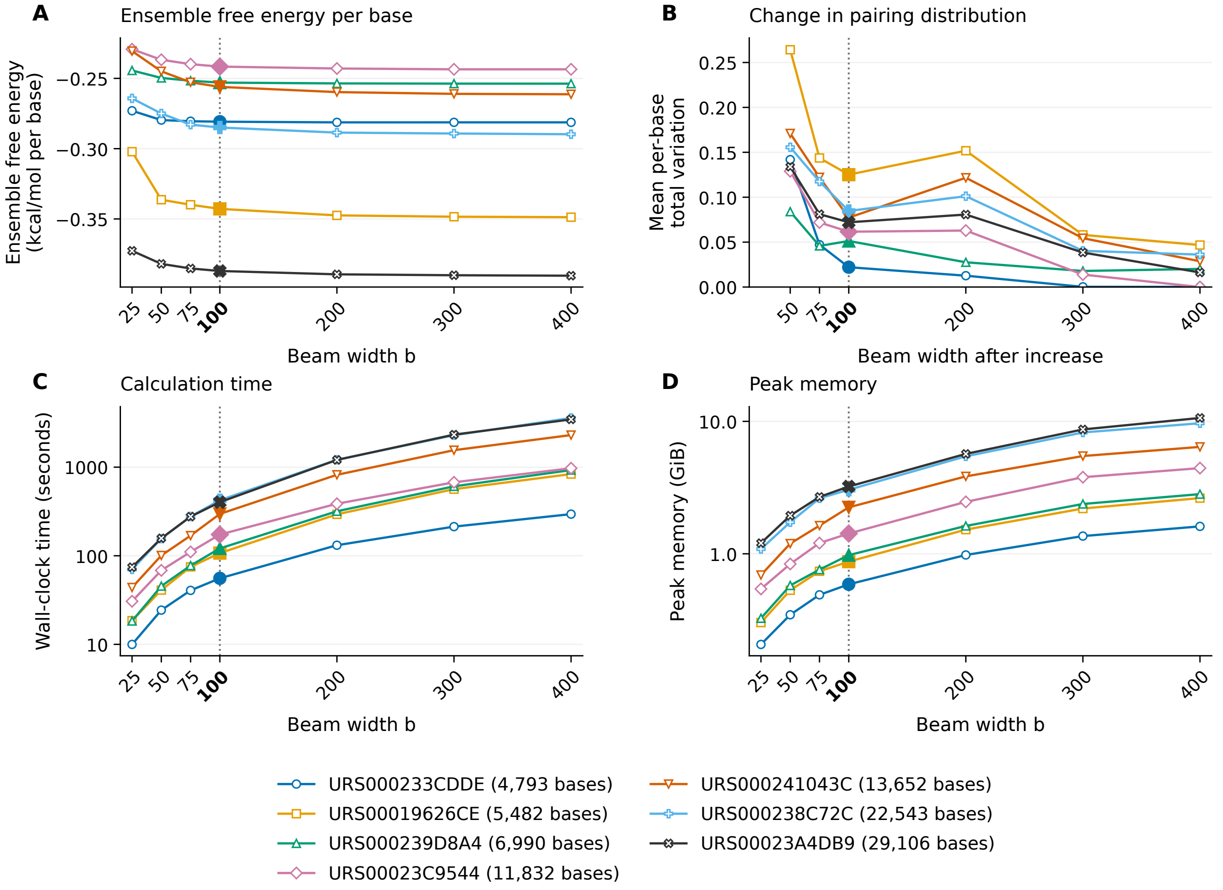


Figure S5: Sensitivity to beam width and local limits, $h = d = \max(b, 30)$, on seven RNAcentral inputs. **(A)** Ensemble free energy per base, $-R_{\text{gas}}T \log \hat{Z}_b/n$ (kcal/mol per base). **(B)** Mean pairing total variation between adjacent widths over bases free in $G$, including the unpaired state. The horizontal coordinate is the width after each increase: $25 \to 50$, $50 \to 75$, $75 \to 100$, $100 \to 200$, $200 \to 300$, and $300 \to 400$. **(C, D)** Whole-process wall time and peak memory, on logarithmic axes. Each line follows one RNA; all 49 settings completed. The dotted guide and larger filled symbols mark $b = 100$. No beam width serves as a reference.

For each RNA, we compared adjacent widths using the mean, p90, and maximum pairing TV over bases free in $G$, including the unpaired state and using probabilities without a cutoff.

Changes generally became smaller as widths increased. For $b = 100 \to 200$, $200 \to 300$, and $300 \to 400$, the maximum absolute free-energy changes per base across the seven RNAs were 0.00463, 0.00126, and 0.000507 kcal/mol per base. The largest pairing TV values, averaged over free bases within each RNA, were 0.1518, 0.0581, and 0.0468 for these three increases, respectively. URS000238C72C (22,543 nt) still had mean TV 0.0361 and maximum per-base TV 0.9991 between $b = 300$ and 400. Changes were not always monotonic. URS000233CDDE's free energy rose by 0.0233 kcal/mol from $b = 200$ to 300. URS000239D8A4's mean TV rose from 0.0179 for $b = 200 \to 300$ to 0.0202 for $300 \to 400$.

At $b = 100$, wall time ranged from 55.7 to 423.5 s and peak memory from 0.588 to 3.222 GiB. At $b = 400$, wall time ranged from 295.0 to 3,592.6 s and peak memory from 1.611 to 10.615 GiB. Each setting used one single-thread process per task on a shared node, with one measurement and no warm-up. Timing covers the partition function, minimum-free-energy calculation, and pairing-probability output, excluding scaffold construction, build, and compression.
